# A Robust Model of Gated Working Memory

**DOI:** 10.1101/589564

**Authors:** Anthony Strock, Xavier Hinaut, Nicolas P. Rougier

## Abstract

Gated working memory is defined as the capacity of holding arbitrary information at any time in order to be used at a later time. Based on electrophysiological recordings, several computational models have tackled the problem using dedicated and explicit mechanisms. We propose instead to consider an implicit mechanism based on a random recurrent neural network. We introduce a robust yet simple reservoir model of gated working memory with instantaneous updates. The model is able to store an arbitrary real value at random time over an extended period of time. The dynamics of the model is a line attractor that learns to exploit reentry and a non-linearity during the training phase using only a few representative values. A deeper study of the model shows that there is actually a large range of hyper parameters for which the results hold (number of neurons, sparsity, global weight scaling, etc.) such that any large enough population, mixing excitatory and inhibitory neurons can quickly learn to realize such gated working memory. In a nutshell, with a minimal set of hypotheses, we show that we can have a robust model of working memory. This suggests this property could be an implicit property of any random population, that can be acquired through learning. Furthermore, considering working memory to be a physically open but functionally closed system, we give account on some counter-intuitive electrophysiological recordings.

## 1 Introduction

The prefrontal cortex (PFC), noteworthy for its highly recurrent connections (Goldman-Rakic, 1987), is involved in many high level capabilities, such as decision making (Bechara et al., 1998), working memory (Goldman-Rakic, 1987), goal directed behavior (Miller and Cohen, 2001), temporal organisation and reasoning (Fuster, 2001). In this article, we are more specifically interested in gated working memory (O’Reilly and Frank, 2006) that is defined as the capacity of holding arbitrary information at a given random time *t*_0_ such as to be accessible at a later random time *t*_1_ (see Figure 1). Between times *t*_0_ and *t*_1_, we make no assumption on the inner mechanisms of the working memory. The only measures we are interested in are the precision of the output (compared to the initial information) and the maximal delay during which this information can be accessed within a given precision range. One obvious and immediate solution to the task is to make an explicit copy (inside the memory) of the information at time *t*_0_ and to hold it unchanged until it is read at time *t*_1_, much like a computer program variable that is first assigned a value in order to be read later. Such solution can be easily characterized by a fixed pattern of sustained activities inside the memory. This is precisely what led researchers to search for such sustained activity inside the frontal cortex (Funahashi, 2017; Constantinidis et al., 2018), where an important part of our working memory capacities is believed to be located. Romo et al. (1999) have shown that PFC neurons of non-human primates can maintain information about a stimulus for several seconds. Their firing rate was correlated with the coding of a specific dimension (frequency) of the stimulus maintained in memory. However, when Machens, Romo, and Brody (2010) later re-analyzed the data of this experiment, they showed that the stimulus was actually encoded over a sub-population using a distributed representation. Similarly, when Rigotti et al. (2013) analyzed single neuron activity recorded in the lateral PFC of monkeys performing complex cognitive tasks, they found several neurons displaying task-related activity. Once they discarded all the neurons that were displaying task-related activity, they were still able to decode task information with a linear decoder. They proposed that the PFC hosts high-dimensional linear and non-linear mixed-selectivity activity. The question is thus, if working memory is not encoded in the sustained activity, what can be the alternatives?

**Figure 1:**
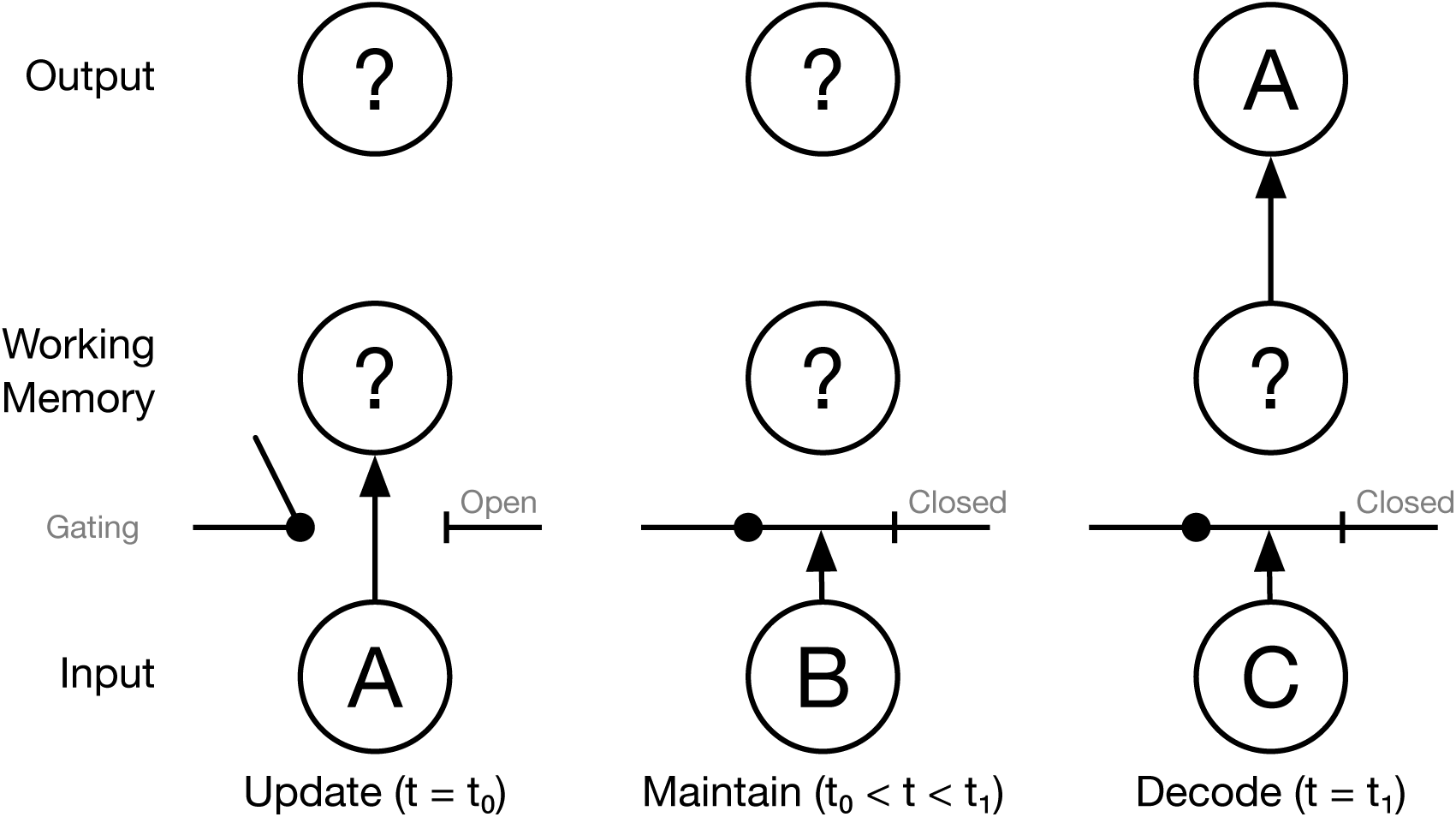
Gated working memory is defined as the capacity of holding arbitrary information at a given random time *t*_0_ such as to be accessible at a later random time *t*_1_. Between times *t*_0_ and *t*_1_, we make no assumption on the activity inside the working memory. Note that a closed gate does not mean external activities cannot enter the memory.

Before answering that question, let us first characterize the type and the properties of information we consider before defining what it means to access the information. The type of information that can be stored inside working memory has been characterized using different cognitive tasks such as for example the delayed matching-to-sample (DMTS), the N-back task or the Wisconsin Card Sorting Task (WCST). From these different tasks, we can assume that virtually any kind of information, be it an auditory or visual stimulus, textual or verbal instruction, implicit or explicit cue, is susceptible to be memorized and processed inside the working memory. From a computational point of view, this can be abstracted into a set of categorical, discrete or continuous values. In this work, we are only interested in the most general case, the continuous one, which can be reduced into a single scalar value. The question we want to address is how a neural population can gate and maintain an arbitrary (e.g. random) value, in spite of noise and distractors, such that it can be decoded non-ambiguously at a later time.

To answer this question we can search the extensive literature on computational models of working memory that have been extensively reviewed by Durstewitz, Seamans, and Sejnowski (2000), by Compte (2006) and more recently by Barak and Tsodyks (2014). More specifically, Compte (2006) explains that the retention of information is often associated with attractors. In the simplest case, a continuous scalar information can be identified with a position on a line attractor. However, this kind of memory exhibits stability issues: unlike a point attractor (i.e. a stable fixed point), a line attractor is marginally stable such that it cannot be robust against all small perturbations. Such a line attractor can be stable against orthogonal perturbations, but not against colinear perturbations (i.e. perturbations along the line). Furthermore, the design (and numerical implementation) of a line attractor is tricky because even small imperfections (e.g. numerical errors) can lead to instability. Nevertheless, there exist several models that can overcome these limitations.

This is the case for the theoretical model by Amari (1977) who proved that a local excitation could persist in the absence of stimuli, in the form of a localized bump of activity in an homogeneous and isotropic neural field model, using long range inhibition and short range excitation. This model represents *de facto* a spatial attractor formed by a collection of localized bumps of activity over the plane. A few decades later, Compte (2000) showed that the same lasting bump property can also be achieved using leaky integrate and fire neurons arranged on a ring, with short range excitation and constant inhibition (ring bump attractor). This model has since then been extended (Edin et al., 2009; Wei, X.-J. Wang, and D.-H. Wang, 2012) with the characterization of the conditions allowing to have simultaneous bumps of activity. This would explain multi-item memorization where each bump represents a different information that is maintained simultaneously with the other bumps. Similarly, Bouchacourt and Buschman (2019) proposed to handle multi-items memorization by duplicating the bump attractor model. They explicitly limited the number of items to be maintained in memory through the interaction between the different bump attractor models (using a random layer of neurons). If all these models can cope with the memorization of a graded information, this information is precisely localized in the bumps of activity and corresponds to a sustained activity. Such patterns of activity have been identified in several cerebral structures (e.g. head direction cells (Zhang, 1996) in mammals, superior colliculus (Gandhi and Katnani, 2011) in primates) but it is not yet clear to what extent this can give account of a general working memory mechanism. Such sustained activity is also present in the model of Koulakov et al. (2002) who consider a population of bistable units that encodes a (quasi) graded information using distributed encoding (percentage of units in high state). This solves both the robustness and stability issue of the line attractor by virtue of discretization. Finally, some authors (Zipser et al., 1993; Lim and Goldman, 2013) consider the encoding of the value to be correlated with the firing rate of a neuron or of a group of neurons. This is the case for the model proposed by Lim and Goldman (2013) who obtain stability of the firing rate by adding a negative derivative self feedback (hence artificially increasing the time constant of neurons). They show how such mechanism can be implemented by the interaction of two populations of excitatory and inhibitory neurons evolving at different time scales. However, independently of the encoding of the graded value, most of model authors are interested in characterizing the mechanism responsible for the maintenance property. They tend to consider the memory as an isolated system, not prone to external perturbations, with the noticeable exception of the model by Zipser et al. (1993) which is constantly fed by an input.

In this work, we consider working memory to be an open system under the constant influence of external activities (i.e. even when the gate is *closed*). Thus, we can not rely on a dynamical system that hosts a line attractor in the absence of inputs. We have to design an input-dependent dynamical system that is robust against all kinds of perturbations (input, internal, output feedback). First, we will formalize a set of tasks that will be used to study features and performances of the different models we have considered. Then, we will introduce a minimal model that will help us in explaining the mechanism needed for a more general model. For this general one, we will consider a particular instance of reservoir: namely an Echo State Network (ESN) (Jaeger, 2001). The analysis of this model will allow us to show that reservoir activity is characterized by a combination of both sustained and transient activities. Moreover, we will show that none of these activities are critical for the decoding of the correct output. Finally, we will show that in the absence of input the dynamics of the model implements a segment attractor (i.e. a line attractor with bounding values).

## 2 Methods

In this section, we formalize and extend the gated working memory task that has been described in the introduction (see Figure 1). We consider four different tasks that will illustrate the generic capacity of the reservoir model to store continuous values or a discrete set of values. The first three are variations of the WM task for continuous values with various number of input values (*n*-value) and gating WM units (*n*-gate). The last task includes a non-linear computation (i.e. digit recognition from pixels) in addition to the gating task for discrete values.

### 2.1 The *n*-value *p*-gate scalar working memory tasks

In the 1-value 1-gate scalar version of the task, we consider at time *t ∈* ℕ an input signal *V* (*t*) in [-1,+1], an input trigger *T* (*t*) in {0, 1} and an output *M* (*t*) that is defined as the value of *V* (*t\**) where *t\** is the most recent time such that *T* (*t\**) = 1 (see Figure 2A). This can be written as:

**Figure 2:**
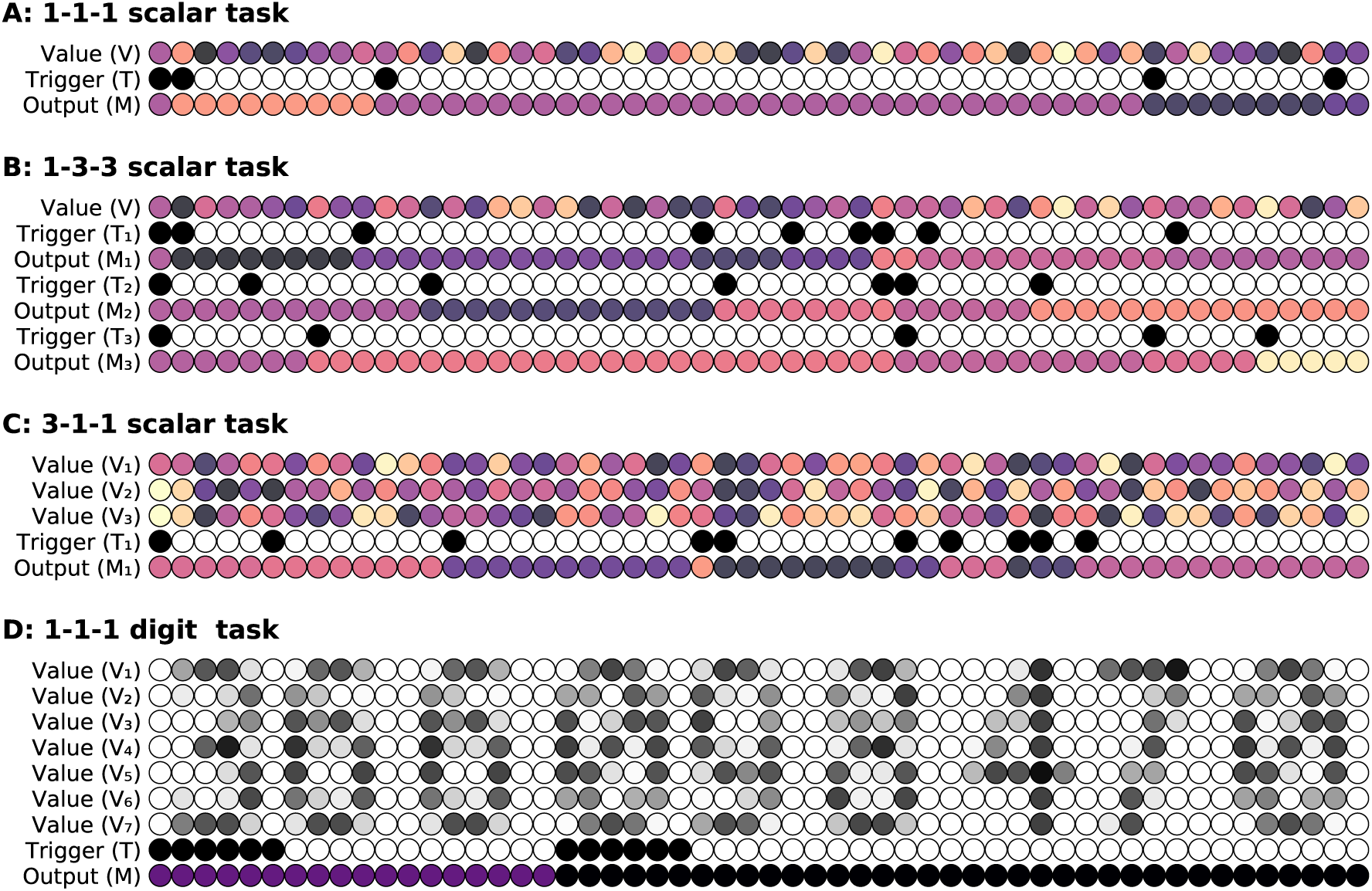
Working memory tasks. Each column represents a time step (time increases from left to right), colored discs represent activity in the input (*V* or *V*_*i*_ and *T* or *T*_*i*_) and the output (*M* or *M*_*i*_). **A.** Gated working memory task with one gate. **B.** Gated working memory task with three gates. **C.** Gated working memory task with one gate but three inputs. To solve the task, it is necessary to ignore the two irrelevant inputs. **D.** Gated working memory task with one gate where the scalar input *V* has been replaced by a streamed input of bitmap digits. To solve the task, it is thus necessary to recognize the digits and to transform them into a normalized value.

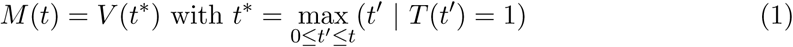

Said differently, the output is the value of the input when the gate was open for the last time. Note that this can be also rewritten as a simple select operator between the input *V* and the output *M* :

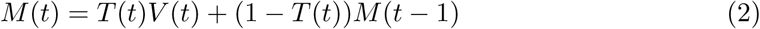

In the 1-value 3-gate version of the task, we consider at time *t ∈* ℕ an input signal *V* (*t*) in [-1,+1], three input triggers *T*_{1,2,3}_(*t*) in 0, 1 and three outputs *M*_{1,2,3}_(*t*) that are respectively defined as the value of *V* (*t\**) where *t\** is the most recent time such that *T*_{1,2,3}_(*t\**) = 1 (see Figure 2B). This can be written as:

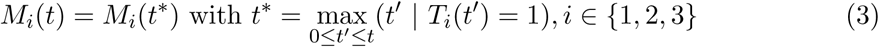

In the 3-value 1-gate scalar version of the task, we consider at time *t ∈* ℕ three input signals *V*_{1,2,3}_(*t*) in [-1,+1], an input trigger *T* (*t*) in {0, 1} and an output *M* (*t*) that is defined as the value of *V*_1_(*t\**) where *t\** is the most recent time such that *T* (*t\**) = 1 (see Figure 2C). This can be written as:

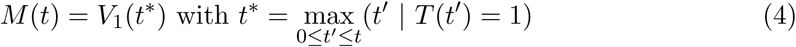

This can be easily generalized to a *n-value p-gate scalar task* with *n* input signals, *p* input triggers, and *p* outputs. Only the first input signal and the *t* input triggers determines the *p* outputs. The other *n -* 1 input signals are additional inputs irrelevant to the task.

### 2.2 The digit 1-value 1-gate working memory task

In the digit version of the 1-value 1-gate working memory task, we define a slightly more complex version of the *V* input by considering a bitmap representation of it (see Figure 2D). Using a monotype font (Inconsolata Regular^1^) at size 11 points, we draw a grayscaled bitmap representation of a sequence of random digits (0 to 9), each digit being of size 6 *×* 7 pixels (after having cropped top and bottom empty lines) and the trigger signal being expanded to the width of a glyph. The output is defined as a discrete and normalized value. It is to be noted that there is no possible linear interpolation between the different inputs, as it was the case for the scalar version. Formally, we can define the output as:

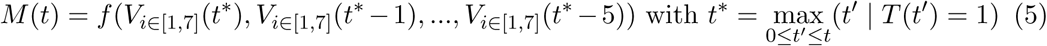

with *f* a function that maps the representation of a digit to a normalized value. Since there are 10 values, the digit *n* is associated to 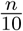. This function has to be learned by the model in order to solve the task. It would have been possible to use the MNIST database (Schaetti, Salomon, and Couturier, 2016) instead of a regular font but this would have also complexified the task, and make the training period much longer, because a digit is processed only when a trigger is present. If we consider for example a sequence of 25,000 digits and a trigger probability of 0.01, this represents (in average) 250 triggers for the whole sequence and consequently only 25 presentations per digit. In comparison, MNIST train data set has 60,000 digits, which would have required as many triggers, and a hundred times more digits.

### 2.3 The minimal model

It is possible to define a minimal degenerated reservoir model (if we consider *X*_1_, *X*_2_ and *X*_3_ to be the reservoir) that takes explicitly advantage of the non-linearity to perform the gating mechanism, as shown in Figure 3. There is no learning in this model. It is parametrized by two values *a* (large enough) and *b* (small enough) and it is fully defined by the following set of equations:

**Figure 3:**
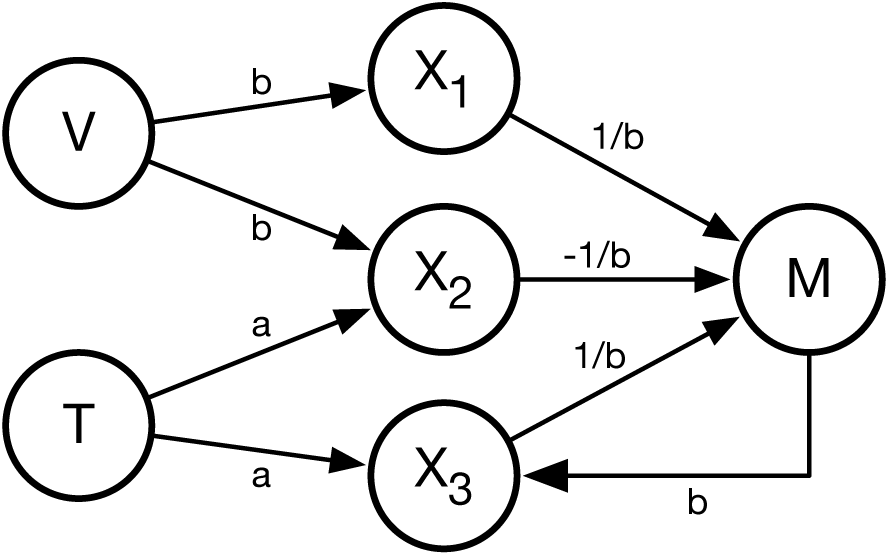
A minimal gated working memory model is able to solve the 1-value 1-gate working memory task by taking advantage of the non-linearity and asymptotic behavior of tanh units. Performance is controlled with parameters *a* and *b*. This architecture can be generalized to the *n-value p-gate task* by adding more than one reservoir unit (corresponding to *X*_3_) for each supplementary trigger/output couple.

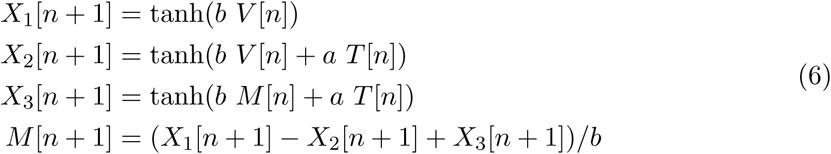

In order to understand how this minimal model works, we can write the output *M* [*n*] relatively to the value of the trigger *T* [*n*] which lies in {0, 1}:

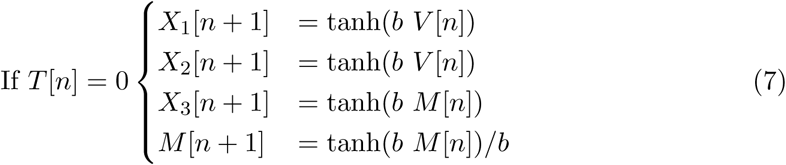

When *T* [*n*] = 0, if we assign *b* a small value (e.g. *b* = 10^*-*3^) and considering that 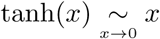, from equation (7), we have *M* [*n* + 1] *≈ M* [*n*].

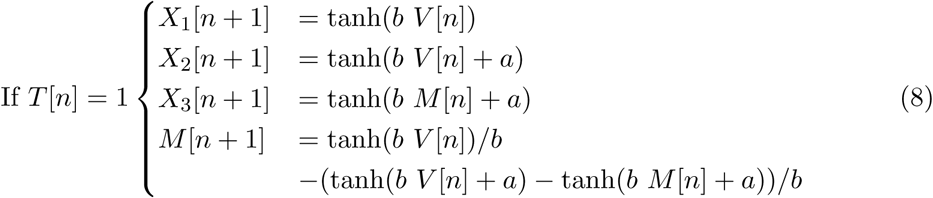

When *T* [*n*] = 1, if we assign *a* to a large value (e.g. *a* = 1000) and considering that *b* is small and that 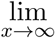 tanh(*x*) = 1, we have *tanh*(*b V* [*n*] + *a*) *≈* tanh(*b M* [*n*] + *a*) *≈* 1. From equation (8), we then have *M* [*n* + 1] *≈* tanh(*b V* [*n*])*/b ≈ V* [*n*].

Consequently, the trigger *T* [*n*] fully dictates the output. When *T* [*n*] = 0, the output *M* [*n* + 1] is unchanged and corresponds to the current memory (*M* [*n*]). When *T* [*n*] = 1, the output *M* [*n* + 1] is assigned the value *V* [*n*] of the input. We think this model represents the minimal model that is able to solve the gating working memory task using such simple neurons (with tanh activation function). By taking advantage of the linear regime around 0 and the asymptotic behavior around infinity, this model gracefully solves the task using only two parameters (a and b). However, we have no formal proof that a model having only two neurons in the reservoir part cannot solve the task.

Note that this minimal model turns out to be similar to the memory cell of a Gated Recurrent Unit (Cho et al., 2014) (GRU) without its reset gate^2^, but using only simple tanh neurons, in comparison to hand-crafted LSTM-like cells. Without the reset gate, the dynamics of a GRU cell can be simplified to:

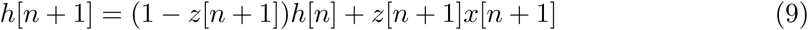

where *h* is the state of the memory cell, *z* is the update gate neuron, and *x* is an input neuron. By using functional equations of hyperbolic functions and classical order 1 Taylor expansions, and for the sake of simplicity by replacing all the *o*_*b→*0_(*b*) by *o*(*b*), we can obtain a similar equation in the minimal model:

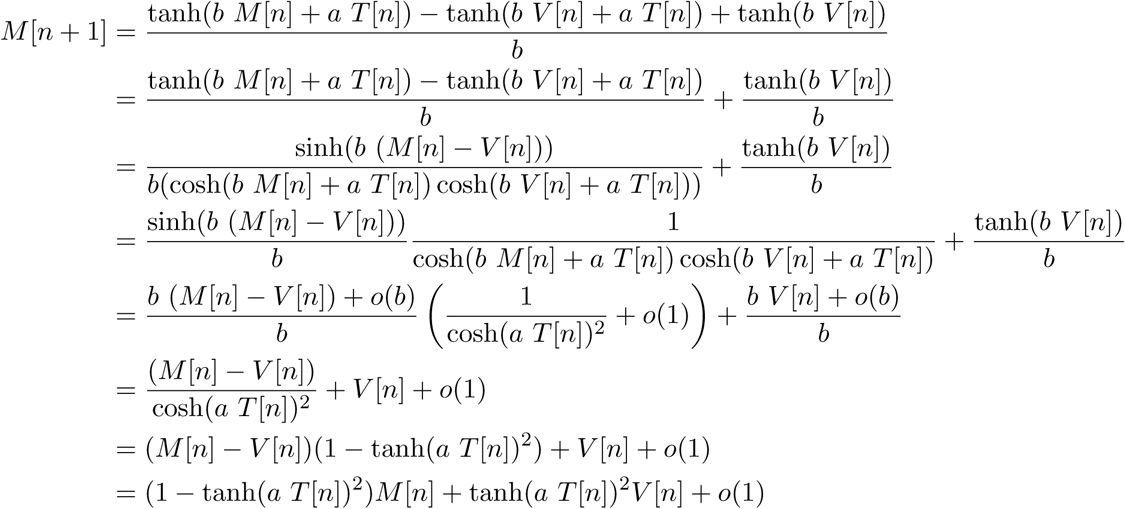

If *b* is small enough, we have *M* [*n* + 1] *≈* (1 *-* tanh(*a T* [*n*])^2^)*M* [*n*] + tanh(*a T* [*n*])^2^*V* [*n*] and consequently, *T* acts as an update gate for *M*. The final form that is equivalent to the GRU without reset gate (as in equation 9) is given by:

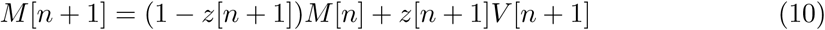

with *z*[*n*] *≈* (1 *-* tanh(*a T* [*n*])^2^).

### 2.4 The reservoir model

We consider an echo state network (Jaeger, 2001) (see Figure 4) with leaky neurons, and feedback from readout units to the reservoir, which is described by the following update equations:

**Figure 4:**
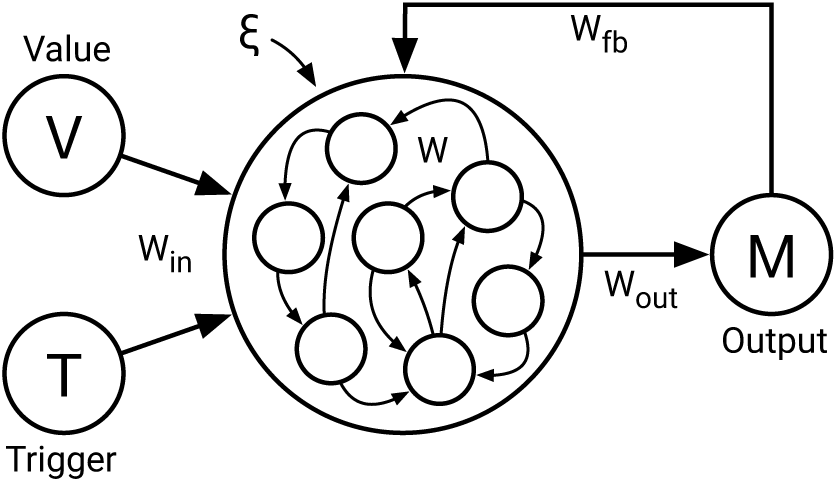
The reservoir model. receives a random signal V in [*-*1, +1] and a trigger signal T in {0, 1}. The reservoir is made of non-linear units (tanh) and the output M is fed back to the reservoir at each iteration. For *n-gate p-value task*, we use *n* triggers, *p* values and *n* outputs. For a generic approach, we use notation *u* for the input, *x* for the reservoir and *y* for the output, independently of the task.

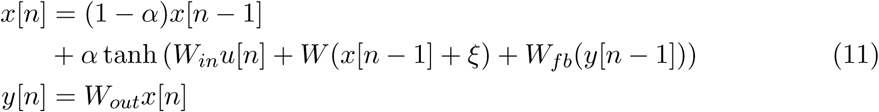

where *u*[*n*], *x*[*n*] and *y*[*n*] are respectively the input, the reservoir and the output at time *n. W*, *W*_*in*_, *W*_*fb*_ and *W*_*out*_ are respectively the recurrent, the input, the feedback and the output weight matrix and *α* is the leaking rate. *ξ* is a uniform white noise term added to the reservoir: *ξ* is independent for each neuron. During initialization, matrices *W*_*in*_ and *W*_*fb*_ are sampled randomly and uniformly between *-*1 and +1 and multiplied by the *input scaling factor* and the *feedback scaling factor* respectively. Matrix *W* is sampled randomly between *-*1 and +1 and we ensure the matrix enforces the defined *sparsity* (ratio of non null values). The resulting matrix is then divided by its largest absolute eigenvalue and multiplied by the specified *spectral radius*. The matrix *W*_*out*_ is initially undefined and is learned through teacher forcing (Lukoševičius, 2012) and a linear regression:

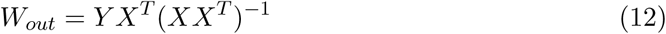

where *Y* and *X* corresponds to the respective concatenation of all the desired outputs and reservoir states during a run, each row corresponding to a time step.

## 3 Results

### 3.1 The reduced model

The reduced model displays a quasi-perfect performance (RMSE = 2e-6 with *a* = 10 and *b* = 10^*-*3^) as shown in Figure 5 and the three neurons *X*1, *X*2 and *X*3 behave as expected. *X*1 is strongly correlated with *V* (Figure 5B), *X*2 is strongly correlated with *V* and saturates in the presence of a tick in *T* (Figure 5C) and *X*3 is strongly correlated with *M* and saturates in the presence of a tick in *T* (Figure 5D). This reduced model is actually a very good approximation of a line attractor (i.e. a line of points with very slow dynamics) even though we can prove that, due to the tanh non-linearity, in the absence of inputs, the model will converge to a null state (possibly after a very long time), independently of parameters *a* and *b* and the initial state. Nonetheless, Figure 5 clearly shows that information can be maintained provided *b* is small enough. There is a drift, but this drift is so small that it can be considered negligible relative to the system time constants: these slow points can be considered as a *line* or *segment attractor* (Seung, 1996; Sussillo and Barak, 2013). As explained by Seung (1998), *the reader should be cautioned that the term “continuous attractor” is an idealization and should not be taken too literally. In real networks, a continuous attractor is only approximated by a manifold in state space along which drift is very slow*.

**Figure 5:**
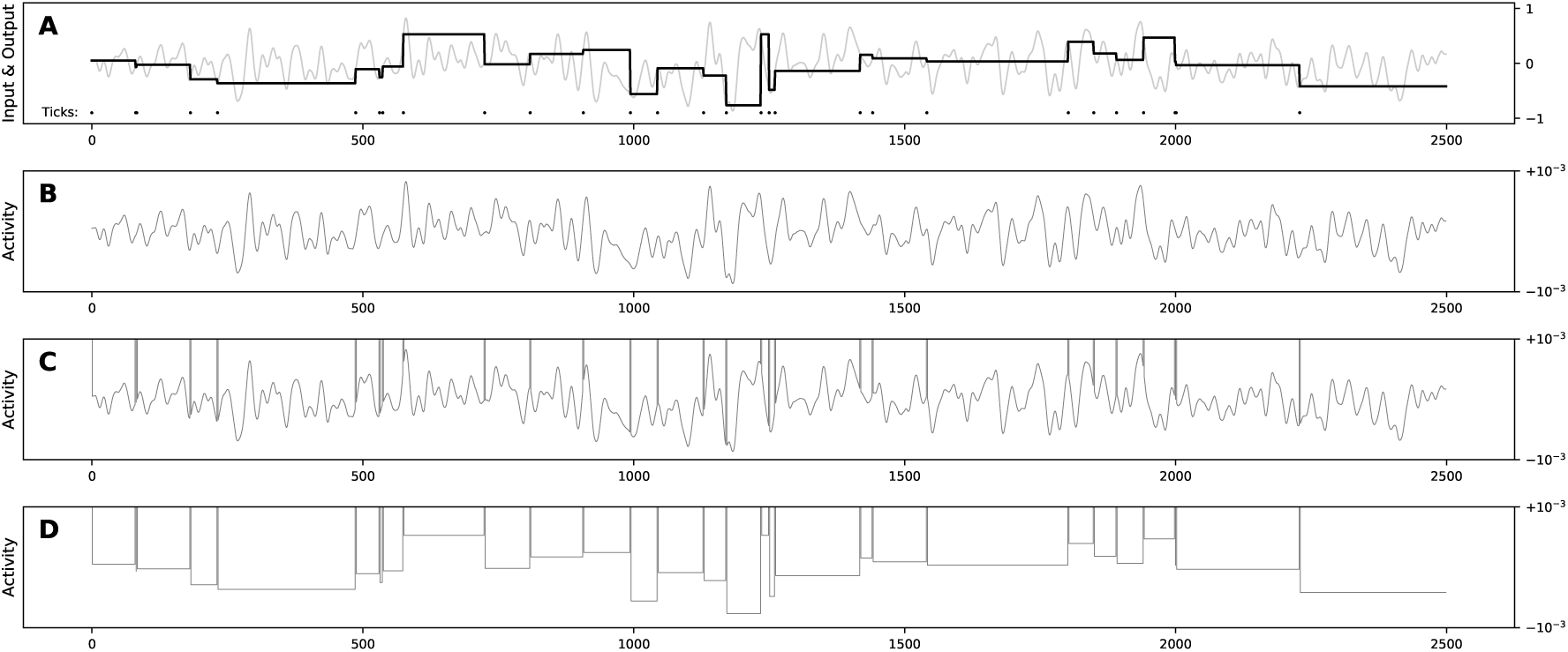
Performance of the reduced model on the 1-gate task. **A** The light gray line is the input signal and the thick black one is the output of the model. Black dots at the bottom represents the trigger signal (when to memorize a new value). **B, C, D** Respective activity of *X*_1_, *X*_2_ and *X*_3_ units.

Nevertheless, it is worth mentioning that in order to have a true line attractor, one can replace the tanh activity function with a linear function saturated to 1 and −1 (Seung, 1996).

### 3.2 The reservoir model

Unless specified otherwise, all the reservoir models were parameterized using values given in table 1. These values were chosen to be simple and do not really impact the performance of the model as it will be explained in the Robustness section. All simulations and figures were produced using the Python scientific stack, namely, SciPy (Jones, Oliphant, and Peterson, 2001), Matplotlib (Hunter, 2007) and NumPy (Walt, Colbert, and Varoquaux, 2011). Sources are available at github.com/rougier/ESN-WM.

**Table 1:** Default parameters. Unless specified otherwise, these are the parameters used in all the simulations.

| Parameter | Value |
| --- | --- |
| Spectral radius | 0.1 |
| Sparsity | 0.5 |
| Leak | 1.0 (no leak) |
| Input scaling | 1.0 |
| Feedback scaling | 1.0 |
| Number of units | 1000 |
| Noise | 0.0001 |
| Training timesteps | 25,000 |
| Testing timesteps | 2,500 |
| Trigger probability | 0.01 |

Results for the reservoir model show a very good generalization performance with a precision of the order of 10^*-*3^ for the level of noise considered (10^*-*4^). Better precision can be obtained for lower noise levels, as show in Figure 7. Surprisingly, this generalization property stands with as few as only four random training values where we can achieve a 10^*-*3^ level of precision.

**Figure 6:**
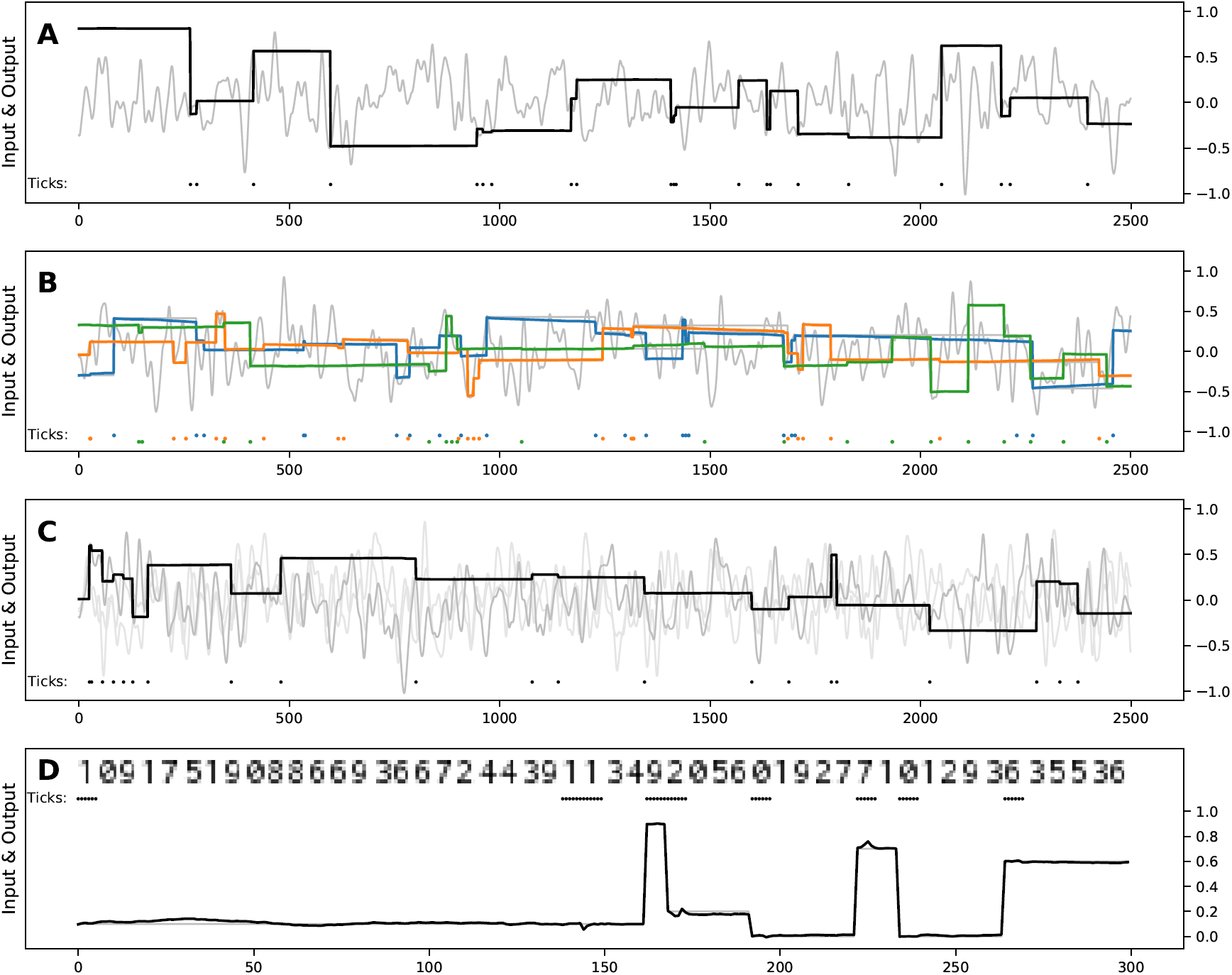
Performance of the reservoir model on working memory tasks. The light gray line is the input signal and the thick black (or colored) one is the output of the model. Dots at the bottom represents the trigger signal (when to memorize a new value). For the digit task, the input containing the value to maintain is shown as a picture instead. **A** *1-value 1-gate scalar task* **B** *1-value 3-gate scalar task* **C** *3-value 1-gate scalar task* **D** *1-value 1-gate digit task*

**Figure 7:**
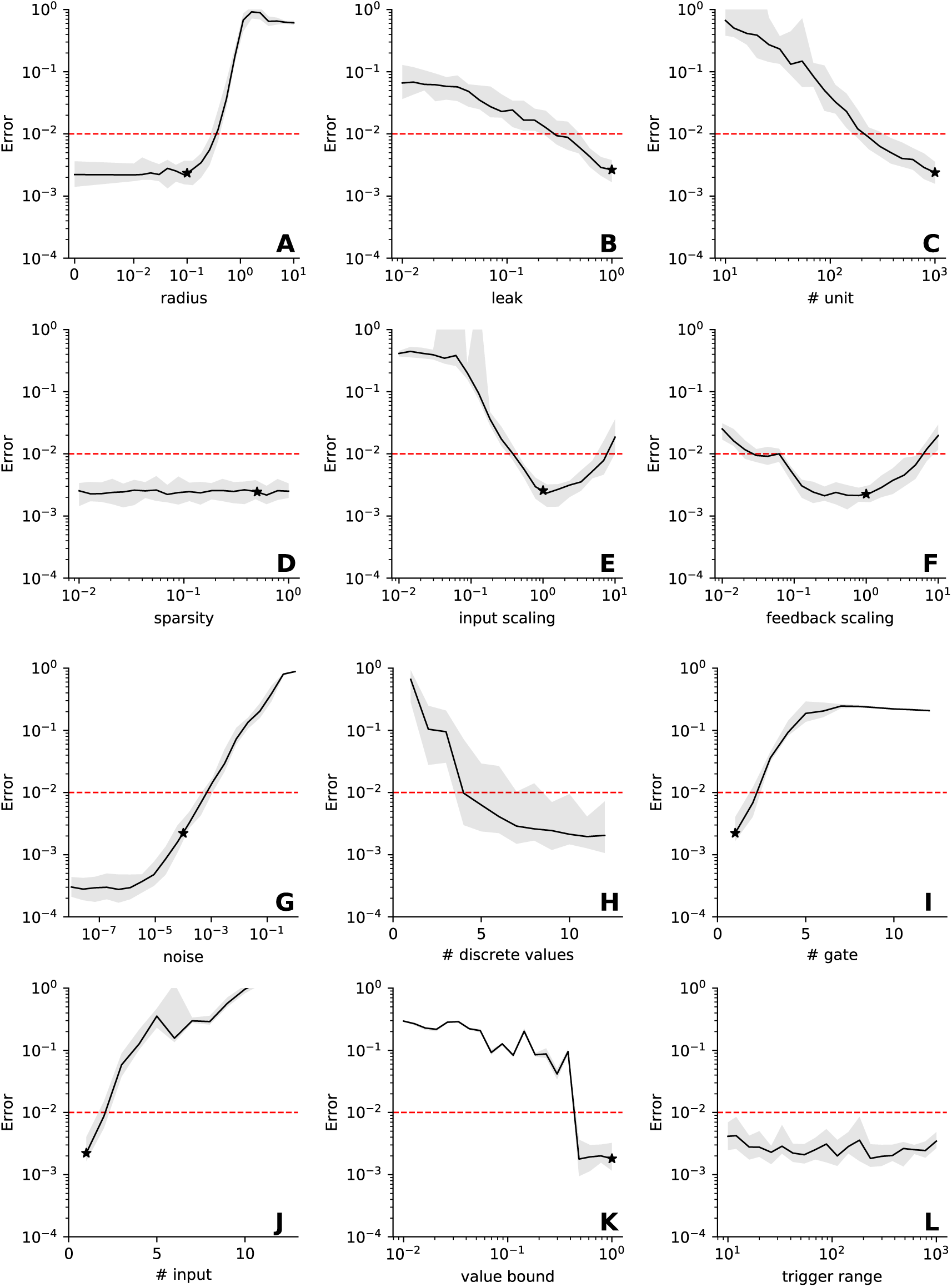
Robustness of the model to hyper-parameters. The performance of the model has been measured when a single hyper-parameter varies while all others are kept constant and equal to the value given in table 1. For each plot, we ran 20 iterations of the model using different random seeds for initialization. Most of the time (**A-GK-L**), the varying parameter has been split into 20 log-uniformly spread values between the minimum and maximum values. **A-F.** Model hyper-parameters. **G-L.** Task hyper-parameters. **A.** Spectral radius ranging from 0.01 to 10. with an additional 0. **B.** Leak term ranging from 0.01 to 1 (no leak). **C.** Number of units in the reservoir (from 1 to 1000). **D.** Sparsity level ranging from 0.01 to 1.0 (fully connected). **E.** Input scaling ranging from 0.01 to 10. **F.** Feedback scaling ranging from 0.01 to 10. **G.** Noise level inside the reservoir, ranging from 10^*-*8^ to 1. **H.** Number of discrete values used to train the model, ranging from 1 to 12. **I.** Number of input gates and output channels, ranging from 1 to 12. (1-value *n*-gate scalar task, RMSE has been divided by the square root of the number of gates) **J.** Number of input values, ranging from 1 to 12. (*n*-value 1-gate scalar task) **K.** Bound for the input value during training, ranging from 0.01 to 1. **L.** Maximal interval (number of steps) between consecutive ticks during training, ranging from 10 to 1000. *\**. Performance for default parameters (Table 1).

#### 3.2.1 1-value 1-gate scalar task

The model has been trained using parameters given in table 1. The *V* signal is made of 25,000 random values sampled from a pseudo-random uniform distribution between −1 and +1. The *T* signal is built from 25,000 random binary values with probability 0.01 of having *T* = 1, and *T* = 0 otherwise. During training, each of the input is presented to the model and the output is forced with the last triggered input. All the input (*u*) and internal (*x*) states are collected and the matrix *W*_*out*_ is computed according to equation (12). The model has been then tested using a *V* signal made of 2500 random values sampled from a pseudo-random uniform distribution between −1 and +1. For readability of the figure (see Figure 6A), this signal has been smoothed using a fixed-size Hann window filter. The corresponding *T* signal has been generated following the same procedure as during the training stage. Figure 6A displays an illustrative test run of the model (for a more thorough analysis of the performance, see the Robustness section) where the error in the output is always kept below 10^*-*2^ and the RMSE is about 3 *\** 10^*-*3^.

#### 3.2.2 1-value 3-gate scalar task

We trained the model on the *1-value 3-gate task* using the same protocol as for the 1-value 1-gate task, using a single value input, three input triggers and three corresponding outputs. Since there are now three feedbacks, we divided respective feedback scaling by 3. Figure 6B shows that maintaining information simultaneously impacts the performance of the model (illustrative test run). There is no catastrophic effect but performances are clearly degraded when compared to the *1-value 1-gate task*. The RMSE on this test run increased by one order of magnitude and is about 2 * 10^*-*2^. Nevertheless, in the majority of the cases we tested, the error does stay below 10^*-*2^. However in a few cases, one memory (and not necessary all) degrades faster.

#### 3.2.3 3-value 1-gate scalar task

We used the same protocol as for the *1-value 1-gate scalar task* but there are now two additional inputs not related to the task and that can be considered as noise. Adding such irrelevant inputs had no effect on the solving of the task as illustrated in Figure 6C that shows an illustrative test run of the model. The error in the output is also always kept below 10^*-*2^ and the RMSE is about the same (3 *\** 10^*-*3^). This is an interesting result, because it means the network is not only able to deal with “background noise” at 10^*-*4^, but it is also able to deal with noise that has the same amplitude as the input. This is an important property to be considered for the modelling of the prefrontal cortex: being an integrative area, the PFC is typically dealing with multimodal information, many of which being not relevant for the working memory task at hand (Mante et al., 2013).

#### 3.2.4 1-value 1-gate digit task

The model has been trained using 25,000 random integer values between 0 and 9 sampled from an uniform distribution. Each of these values is drawn one after the other onto an image, each character covering 6 columns of the image. The input *V* consists then of the row of this image. The *T* signal is sampled similarly as in the *1-value 1-gate task* and then expanded 6 times to fit the transformation of the value to the picture of the values. Which means that trigger lasts 6 time steps. Interestingly, as we show on Figure 6D, even if the value to maintain is not explicit anymore, it can still be extracted and maintained. On the test run we show the RMSE is about 4 *\** 10^*-*2^. It is to be noted that the recognition of a digit is not straightforward and may require a few timesteps before the digit is actually identified. However when the good value is maintained it seems to last, the absolute error stays below 0.05, which is the threshold from which we can distinguish between two values. The reservoir parameters that we found are robust enough to enable not only a pure memory task (i.e. gating), but also a discrimination task (i.e. digit recognition).

Dambre et al. (2012) demonstrated the existence of a universal trade-off between the non-linearity of the computation and the short-term memory in the information processing capacity of any dynamical systems, including echo state networks. In other words, the hyperparameters used to generate an optimal reservoir for solving a given memory task would not be optimal for a non-linear computation task. Here we see that even if the reservoir is made to memorised a particular value, it is still able to do a non-linear task such as discriminating stream of digits. Pascanu and Jaeger (2011) initiated the concept of such working memory (WM) units for reservoirs. They processed streams of characters using six binary WM-units to record the deepness of curly brackets that appeared. We made such reservoir-WM-units coupling more general: from binary to continuous values. Instead of relying on a collection of N binary WM-units to encode N values, or on the maximum encoding of 2^*N*^ values, we have shown that a reservoir can use only one WM-unit to encode a continuous value with a good precision.

### 3.3 Robustness

We analyzed the robustness of the model first by measuring its sensitivity for each of the hyper-parameters (Figure 7A-F), namely: input scaling, feedback scaling, spectral radius, sparsity of the reservoir weight matrix, leak term (*α*) in equation (11) and number of units in the reservoir. We also we measured its sensitivity to task hyper-parameters (Figure 7G-L): the noise level (*ξ*), number of discrete values used during training (when there is a trigger, *V* is sampled uniformly in uniformly sampled between −1 and 1 discrete values), the temporal delay between successive gating signals in training (*T* is built sampling its interval between triggers uniformly between 0 and a bound), the bound used to sample the input value in training (*V* is uniformly sampled between -*b* and *b* instead, where *b* is the bound), the total number of input gates (with a corresponding number of outputs), the number of input values. For most hyper-parameter, we set a minimum and maximum value and pick 20 values logarithmically spread inside this range. For each task and model hyperparater we ran 20 simulation instances for 25,000 timesteps and record the mean performance using 2,500 values. Results are shown in Figure 7.

First, we can see a non-sensitivity to the sparsity (i.e. minor differences in performances when these parameters vary). Similarly we can see a non-sensitivity to the leak term, input and feedback scaling, as long as they are not too small. It is to be noted that input and feedback scaling should also not be too big. As expected, the performance increases with the number of neurons. Surprisingly, we can note that the performance decreases with the spectral radius. In fact, in supplementary Figure 17 we analysed the behavior of the reservoir model with various spectral radii. We show that even with a bigger spectral radius, the reservoir keeps maintaining something relevant but less precise (the segment attractor is slowly degenerating). Globally, the reservoir model is very robust against model hyper-parameters changes as long as it is trained in this condition.

Concerning the task hyper-parameters, one can see in Figure 7 that only the trigger range has no impact. This means that, whatever the time elapsed between two triggers, it does not affect the performance. Performance naturally decreases with the increase of the noise level (Figure 7**G**), the number of input gates (**I**) or the number of input values (**J**). We can note that the number of discrete values used during training impacts the performance in a very specific way (**H**). Using between 4 to 7 training values, the performance is already good and does not improve further with supplementary training values. This means that even if the reservoir model has been only trained to build few stable points, it is able to interpolate and maintain the other points on the segment attractor. Interestingly, in Figure 7**K**, we can see a similar case of interpolation relatively to the input value bound *x* (i.e. the interval [*-x, x*] on which the output is trained). The performance reached a plateau when the bound values reached 0.5 while the interval used for testing performance is always [*-*1, 1].

### 3.4 Dynamics

#### A segment attractor

Figure 8 shows how the model evolves after having been trained for the *1-value 1-gate task* using different starting positions and receiving no input. This results in the formation of a segment attractor even tough the model was only trained to gate and memorize continuous values. If we compute the principal component analysis (PCA) on the reservoir states and look at the principal components (PCs) in the absence of inputs, we can observe (supplementary Figure 16) that all the reservoir states are organized along a straight line on the first component (the one which explains most of the variance) and each point of this line can be associated with a corresponding memory value. Interestingly enough, there are points on this line that correspond to values outside the [*-*1; 1] range, i.e. values for which the model has not been trained for. However, these points are not stable and an any dynamics starting from these points converge towards the points associated to the values 1 or −1 (see Figure 8).

**Figure 8:**
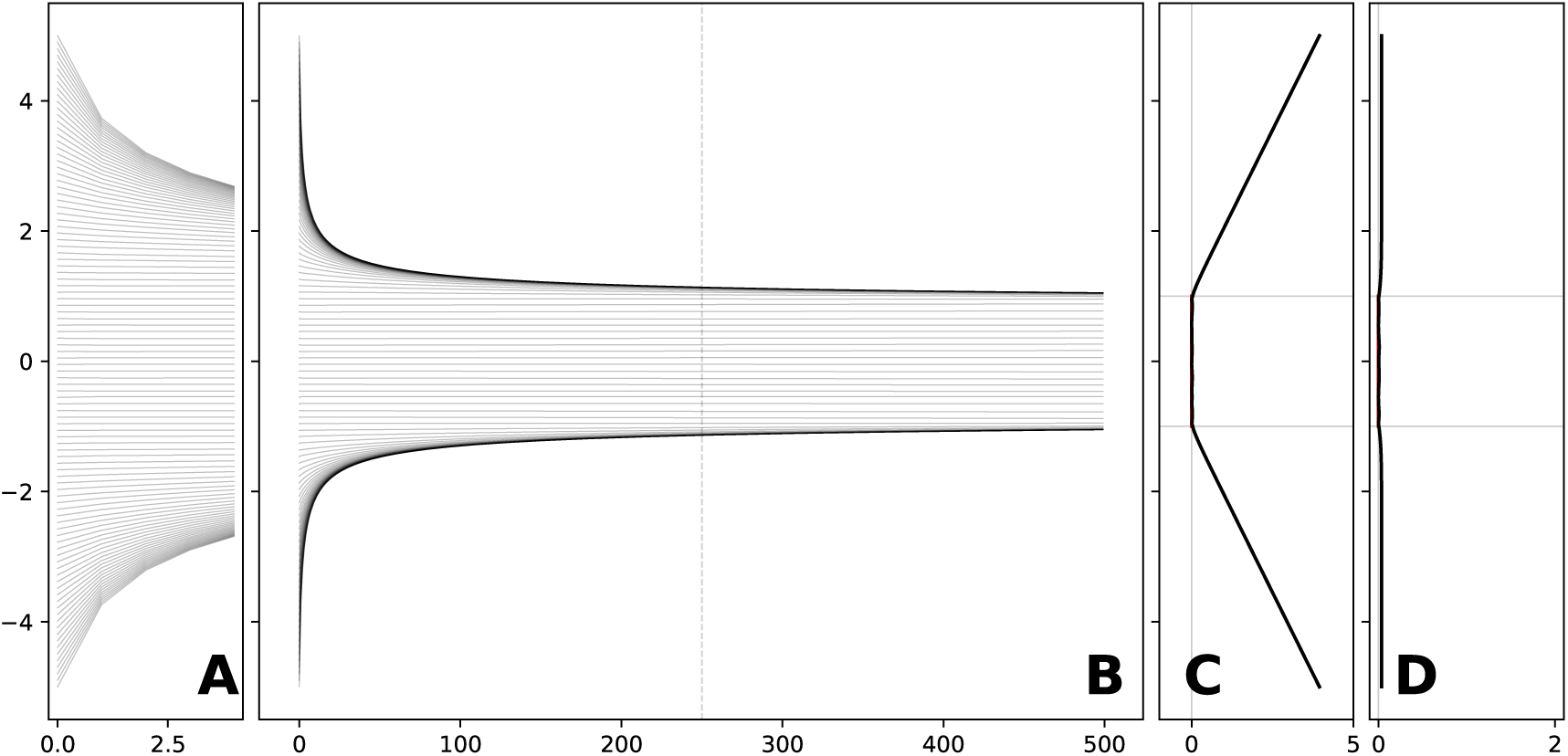
An approximate segment attractor. The same model trained on the 1-value 1-gate was tested for 500 iterations without inputs, starting with an initial trigger along with values linearly distributed between −5 and +5. **A** and **B** Output trajectories. Each line corresponds to one trajectory (output value) of the model. **A** is a zoom of **B** on the first few time steps. **C** Measure of how well are maintained the initial triggered value: the absolute difference in the output between initial time and final time. **D** Measure of how stable are the states reached by the reservoir: The maximal absolute difference between states at intermediate time (dashed line) and final time.

#### *V*-like and *M*-like neurons

Similarly to the minimal model, in the absence of input, the inner dynamic of the reservoir model is a combination of both sustained and highly variable activities. More precisely, in the *1-value 1-gate task* we notice two types of neurons that are similar to neurons *X*1 and *X*3 in the reduced model: (1) neurons which solely follows the input *V* (i.e. *V*-like neurons), and (2) neurons that mostly follow the output *M* (i.e. *M*-like neurons), respectively. We also notice that *M* (*|w| ≤* 0.1)-like neurons average activity is linearly linked with the *M* value and fluctuate around this mean activity according to the input *V*. In Figure 9 we show *M*-like neurons for the different tasks. These neurons were found by taking the most correlated neurons with the output M^3^. From Figure 9**A** to **D** we see that the M-like neurons link with *M* output goes weaker and weaker: the average sustained activity goes from nearly flat to highly perturbed.

**Figure 9:**
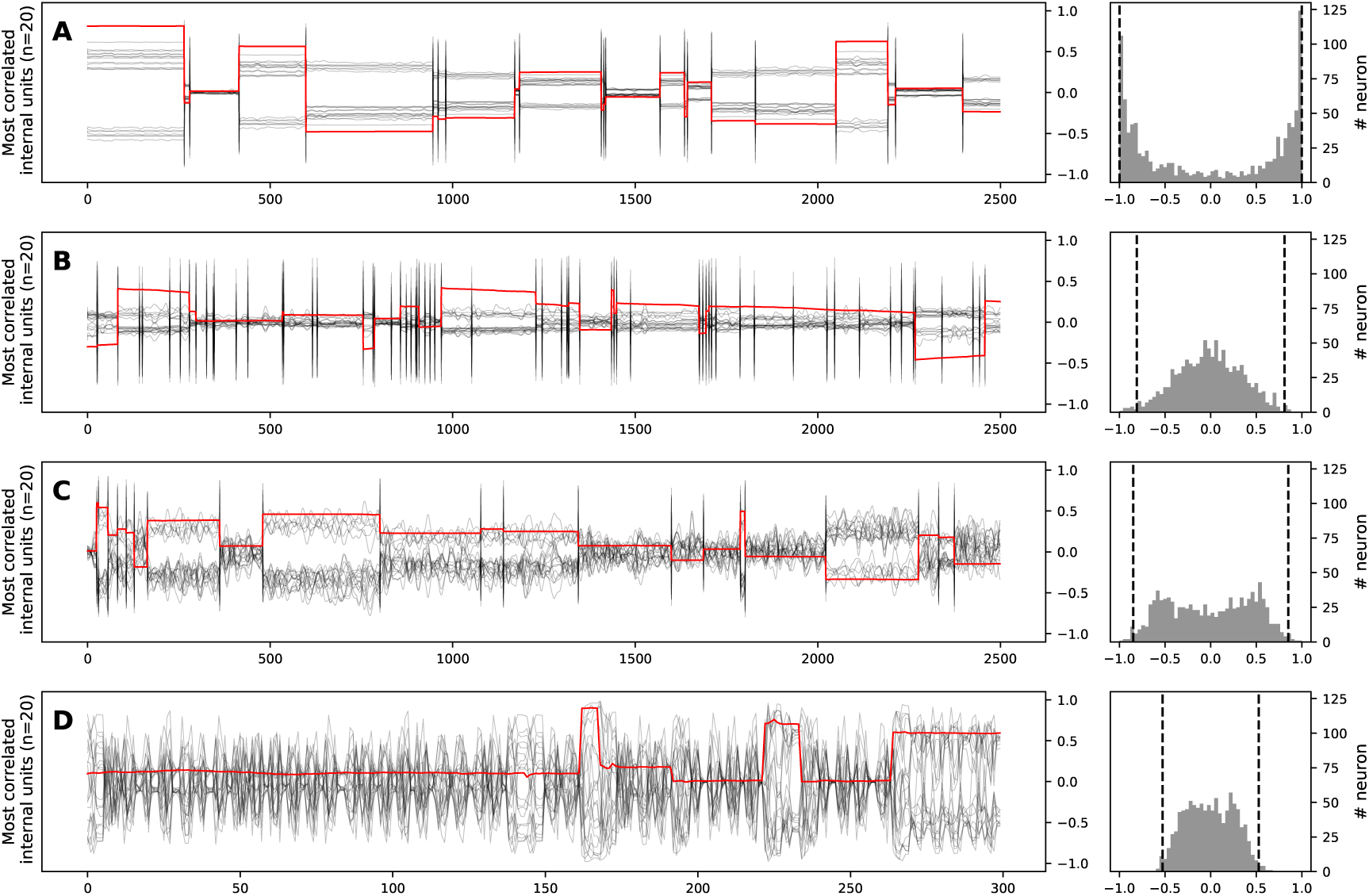
Most correlated reservoir units displaying various degrees of maintained activity. Left: in black the activities of the 20 neurons the most correlated with the output, in red the output of the model. Right: histogram of the correlation between the neurons and the output; y-axis represents the number of neurons correlated with the output produced by the model within a correlation bin. Dashed lines indicate the 20th most correlated and most anti-correlatted neurons with the output. **A** *1-value 1-gate scalar task* **B** *1-value 3-gate scalar task* **C** *3-value 1-gate scalar task* **D** *1-value 1-gate digit task*. Most correlated reservoir units do not necessarily show clear sustained activity: we can see a degradation of sustained activities from **A** to **D** according to the different tasks. Thus, sustained activity is not mandatory to perform working memory tasks.

This “degradation” of sustained activity is explained by the change in the distribution of correlations of the whole reservoir population with *M* output: in Figure 9 **(right)** we see that the correlation with *M* output is quickly shrinking from **A** to **D**. For the *1-value 1-gate task* (9A) mostly all neurons stay at the same value while maintaining the memory. However, when more values have to be maintained (Figure 9B) or when more inputs are received (Figure 9C), most of the activities do not stay at the same value anymore while maintaining the memory. In fact, in the *1-value 1-gate digit task*, neurons do not display a sustained activity at all (Figure 9D). Interestingly, similar behavior (no sustained activity) can be obtained by lowering the feedback scaling (supplementary Figure 18) or by enforcing the input weights to be big enough (supplementary Figure 20). More formally, when there is no trigger, the activity of the neurons can be rewritten as tanh(*aX* + *bM*). The two proposed modifications make the ratio 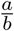 bigger and eventually when *a* ≫ *b*, tanh(*aX* + *bM*) *≈* tanh(*aX*). Consequently, tanh(*aX*) is highly correlated with *X* as *aX* stays bounded between −1 and 1, and does not depend of *M*. Similarly, when *a ≪ b*, tanh(*aX* + *bM*) *≈* tanh(*bM*) which is in turn highly correlated with *M* for the same reasons.

#### Linear decoder

To go further in the understanding of the role played by sustained activities, we wanted to know how much of these sustained activities were necessary to decode the output memory *M*. For the *1-value 1-gate task*, we trained a separate decoder based on a subsample of the reservoir population. We increasingly subsampled neurons based on three conditions: either by choosing the most correlated one first, the least correlated one first or just randomly selecting them. In Figure 10 we can see two interesting facts. First, there is no need of the whole reservoir population to decode well enough the memory *M*: taking a hundred neurons among one thousand is sufficient. Second, if we take enough neurons, there is no advantage in taking the most correlated one first, random ones are enough. Surprisingly, it seems better to rely on randomly selected units than most correlated ones. This suggests that randomly distributed activities contained more information than just most correlated unit activities and offers a complementary perspective when comparing it to Rigotti et al. (2013) decoding of neural population recorded in monkeys: when task-related neurons are not kept for decoding, information (but less accurate) can still be decoded.

**Figure 10:**
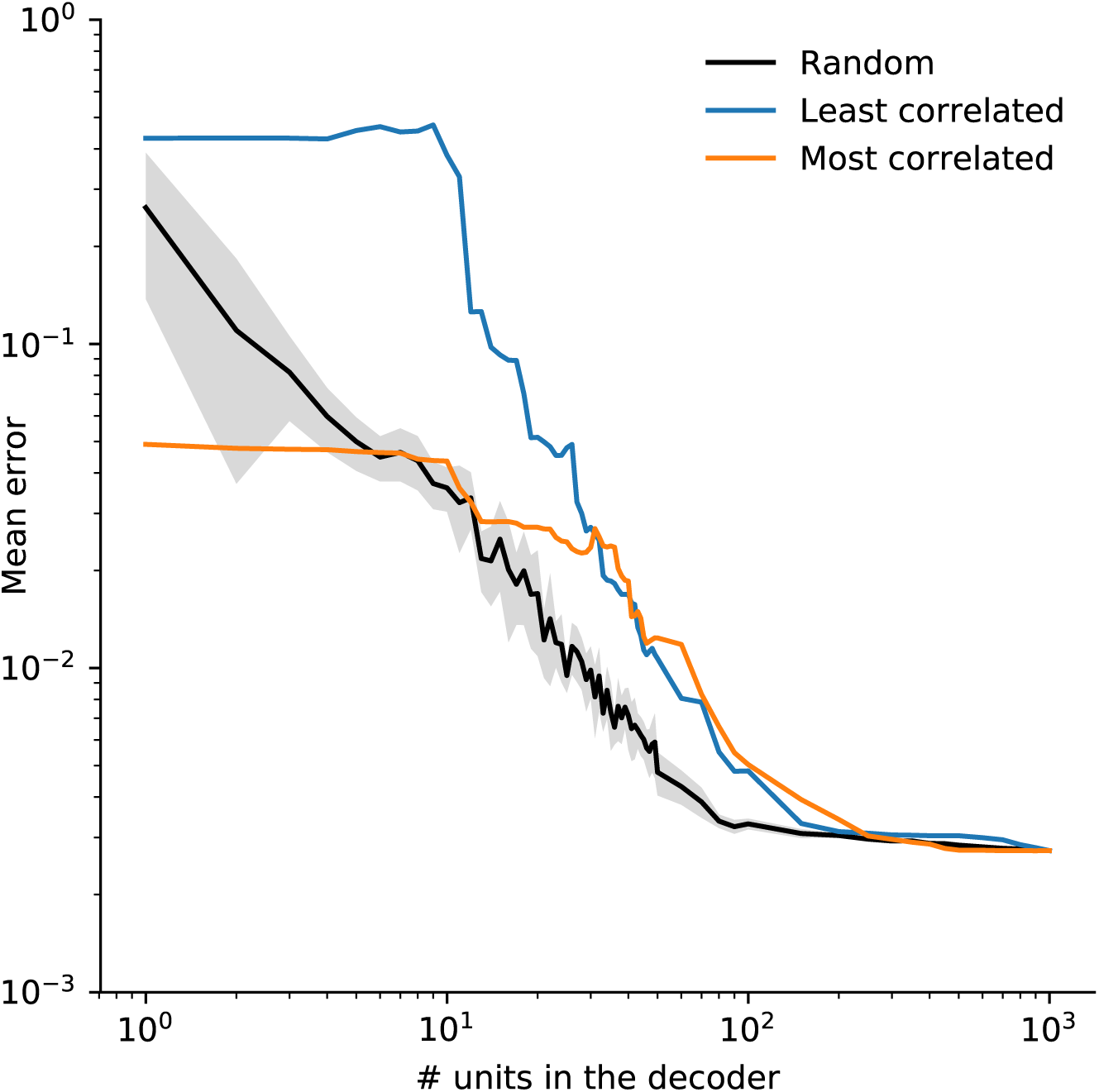
Performance of variable size linear decoders. After training, we ran the model on the training set and recorded all the internal states. We selected a subset (from 1 to 1000 units) of these internal states and find the associated best linear decoder on the training data set. Finally, we measured the performance of these decoders on the training sets. Units composing a subset have been sampled either using the least or the most correlated units (with the output) or simply randomly. Performance for subsets of size 1000 is equal to the performance of the original model.

### 3.5 Equivalence between the minimal and the reservoir model

In order to understand the equivalence between the minimal and the reservoir model, it is important to note that there are actually two different regimes as shown in Figure 11. One regime corresponds to the absence of a trigger (T=0) and the other regime corresponds to the presence of a trigger (T=1). When there is no trigger (T=0), the activities of *X*_1_ and *X*_2_ compensate each other because they are in quasi-linear regime (*b* being very small) and their summed contribution to the output is nil.

**Figure 11:**
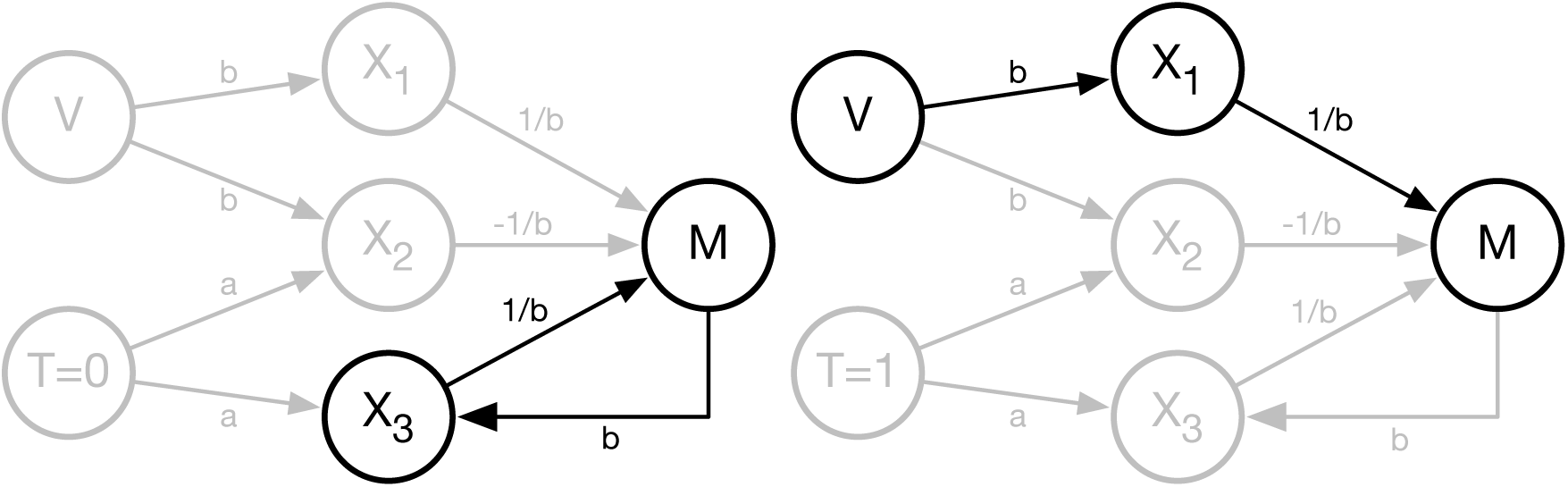
The two regimes of the minimal model depending on the absence (right) or the presence (left) of a trigger T (0 or 1).

In the reservoir model, we can actually identify an equivalent population by discriminating neurons inside the reservoir based on the strength of their input weight relatively to *V*. More formally, we define *R*_12_ as *the group of neurons whose input weight from V (absolute value) is greater than* 0.1. Figure 12 (panel H) shows that the summed output of these neurons is quasi nil while the complementary population *R*_3_ is fully correlated with the output and is thus equivalent to the *X*_3_ unit in the minimal model.

**Figure 12:**
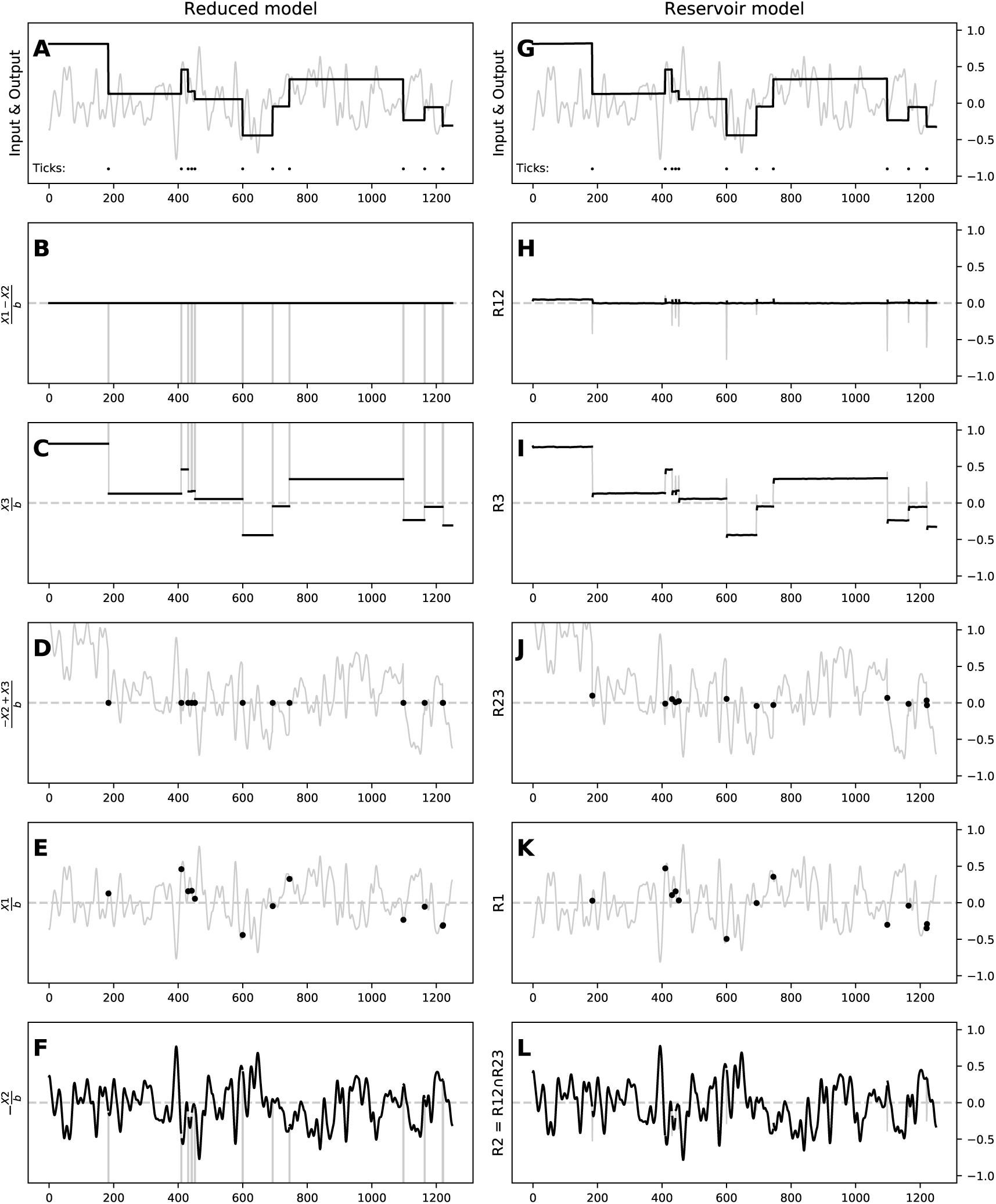
Side by side comparison of the minimal (left) and full (right) models. **A & G** Output of the two models respectively. **B & H** Group of neurons whose sum of activity is nil between triggers. **C & I** Group of neurons whose sum of activity is equal to the output between triggers. **D & J** Group of neurons whose sum of activity is nil during triggers. **E & K** Group of neurons whose sum of activity is equal to the input during triggers. **F & L** Group of neurons whose sum of activity is equal to the input between triggers.

Symetrically, in the presence of a trigger (T=1), the activities of *X*_2_ and *X*_3_ compensate each other because they are in saturated regime (*a* being very large) and their summed contribution to the output is nil. We can again identify an equivalent population in the reservoir model by discriminating neurons inside the reservoir based on the strength of their input weight relatively to *T*. More formally, we define *R*_23_ as *the group of neurons whose input weight from T (absolute value) is greater than* 0.05. Figure 12 (panel J) shows that the summed output of these neurons is quasi nil while the complementary population *R*_1_ is fully correlated with the input *V* is thus equivalent to the *X*_1_ unit in the minimal model.

We can then identify *R*_2_ by taking the intersection of *R*_12_ and *R*_23_ whose activity is similar to *X*_3_ (see panel L in Figure 12). Consequently, we have identified in the reservoir non-disjoint sub-populations that are respectively and collectively equivalent to activity of *X*_1_, *X*_2_ and *X*_3_ in the minimal model.

## 4 Discussion

In computational neuroscience, the reservoir computing paradigm (RC) (Jaeger, 2001; Maass, Natschläger, and Markram, 2002; Verstraeten et al., 2007), originally proposed independently by Dominey (1995), and Buonomano and Merzenich (1995)^4^, is often used as a model of canonical microcircuits (Maass, Natschläger, and Markram, 2002; Hoerzer, Legenstein, and Maass, 2012; Sussillo, 2014). It is composed of a random recurrent neural network (i.e. a reservoir) from which readout units (i.e. outputs) are trained to linearly extract information from the high-dimensional non-linear dynamics of the reservoir. Several authors have taken advantage of this paradigm to model cortical areas such as PFC (Hinaut and Dominey, 2013; Mannella and Baldassarre, 2015; Hinaut, Lance, et al., 2015; Enel et al., 2016) because most of the connections are not trained, especially the recurrrent ones. Another reason to use the reservoir computing paradigm for PFC modelling is because PFC also hosts high-dimensional non-linear dynamics (Rigotti et al., 2013). RC offers a neuroanatomically plausible view of how cortex-to-basal-ganglia (i.e. cortico-basal) connections could be trained with dopamine: the reservoir plays the role of a cortical area (e.g. trained with unsupervised learning), and the read-out units play the role of basal ganglia input (i.e. striatum).

However, in many dynamical systems, reservoirs included, there exists a trade-off between memory capacity and non-linear computation (Dambre et al., 2012)^5^. This is why some studies have focused on reservoirs with dedicated readout units acting as working memory (WM) units (Hoerzer, Legenstein, and Maass, 2012; Pascanu and Jaeger, 2011; Nachstedt and Tetzlaff, 2017). These WM units have feedback connections projecting to the reservoir and are trained to store binary values that are input-dependent. This somehow simplifies the task and enables the reservoir to access and use such long-time dependency information to perform a more complex task, freeing the system from constraining reservoir short-term dynamics. Such ideas had already some theoretical support, for instance Maass, Joshi, and Sontag (2007) showed that with an appropriate readout and feedback fonctions, readout units could be used to approximate any k-order differential equation. Pascanu and Jaeger (2011) used up to six binary WM units to store information in order to solve a nested bracketing levels task. Using a principal component analysis, they showed that these binary WM units constrain the reservoir in lower dimensional “attractors”. Additionally, Hoerzer, Legenstein, and Maass (2012) showed that analog WM units (encoding binary information) also drive the reservoir into a lower dimensional space (i.e. 99% of the variability of the reservoir activities are explained by fewer principal components). More recently, Strock, Rougier, and Hinaut (2018), Beer and Barak (2019) used such WM units in order to store analog values (as opposed to binary ones) in order to build a *line attractor* (Seung, 1996; Sussillo and Barak, 2013). In particular, Beer and Barak (2019) explored how a line attractor can be built online, by comparing FORCE (Sussillo and Abbott, 2009) and LMS algorithms, using a WM unit to maintain a continuous value in *the absence* of input perturbations.

In that context, the minimal model of three neurons we have proposed helps to understand the mechanisms that allows a reservoir model to gate and maintain scalar values, in *the presence* of input perturbations, instead of just binary values. As explained previously, this minimal model exploits the non-linearity and the asymptotic behavior of the three tanh units and mimics a select operator between the input signal and the output. In the case of the reservoir model, there is no precise architecture or crafted weights, but this is compensated by the size of the population inside the reservoir along with the training of the output weights. More precisely, we have shown that virtually any population of randomly connected units is able to maintain an analog value at any time and for an arbitrary delay. Taking advantage of the non-linearity of the neuron transfer function, we have shown how such population can learn a set of weights during the training phase, using only a few representative values. Given the random nature of the architecture and the large set of hyper-parameters, for which the precision of the output remains acceptable, this suggests that this property could be a structural property of any population that could be acquired through learning. To achieve such property we mainly used offline training in our analyses (for efficiency reasons), but we have shown that it also works with online FORCE learning (see supplementary materials and Figure 14 and 15).

We have shown that the reservoir model behavior is similar to the minimal model with the presence of two “macro states” that are implemented by compensatory clusters. In a nusthell, this working memory is using two distinct mechanisms: a selection mechanism (i.e. a switch), and a line attractor. Such mechanims have been also reported in a fully trained recurrent neural network with back-propagation (Mante et al., 2013). Authors proposed a context-dependent selective integration task and shows that “the rich dynamics of PFC responses during selection and integration of inputs can be characterized and understood with just two features of a dynamical system — the line attractor and the selection vector, which are defined only at the level of the neural population”. However in our case, not only did we rely on the analysis of the dynamical system to understand the behavior of the system, but we were able to design a minimal model implementing these mechanisms and show these same mechanisms are also present in the reservoir model but in a distributed way.

Finally, one important feature of the model is that it is actually an open system and as such, it is continuously under the direct influence of external activities. More precisely, the model is able to retain an information when the gate is closed, but this *closed gate* corresponds to a functionnal state rather than a physical state where input signals would have been blocked and information continues to enter the system. This is illustrated quite clearly when one looks at the internal activities inside the reservoir: a large number of neurons are directly (and only) correlated with the input signals. This has consequences for the analysis of the dynamics of the population: this population is partly driven by the (working memory) task and partly driven by uncorrelated external activities. If we go back to biology, this makes perfect sense: a population of neurons is never isolated from the rest of the brain. When studying population electrophysiological recordings, we have to keep in mind that such activities can be fully uncorrelated with the task we are observing. This might be one of the reasons for the variability of hypotheses about working memory encoding mechanisms.

## 5 Supplementary

### 5.1 Reduced model

The effect of parameters *a* and *b* are of distinct nature as illustrated in Figure 13. *b* controls the time the memory takes to fade to zero whereas *a* controls how the model will mix the previous output and the new desired one. Interestingly, the time the information takes to fade out to zero is not directly linked to a time constant of neurons. Moreover, the mixing factor can actually be computed for *b* small enough and is equal to tanh(*a*)^2^. Consequently, by feeding *T* with *a* (instead of 1) and fixing the input weights coming from *T* to 1 (instead of *a*), one can obtain a model which mimics the update mechanism of a Gated Recurrent Unit (GRU) but with regular neurons.

**Figure 13:**
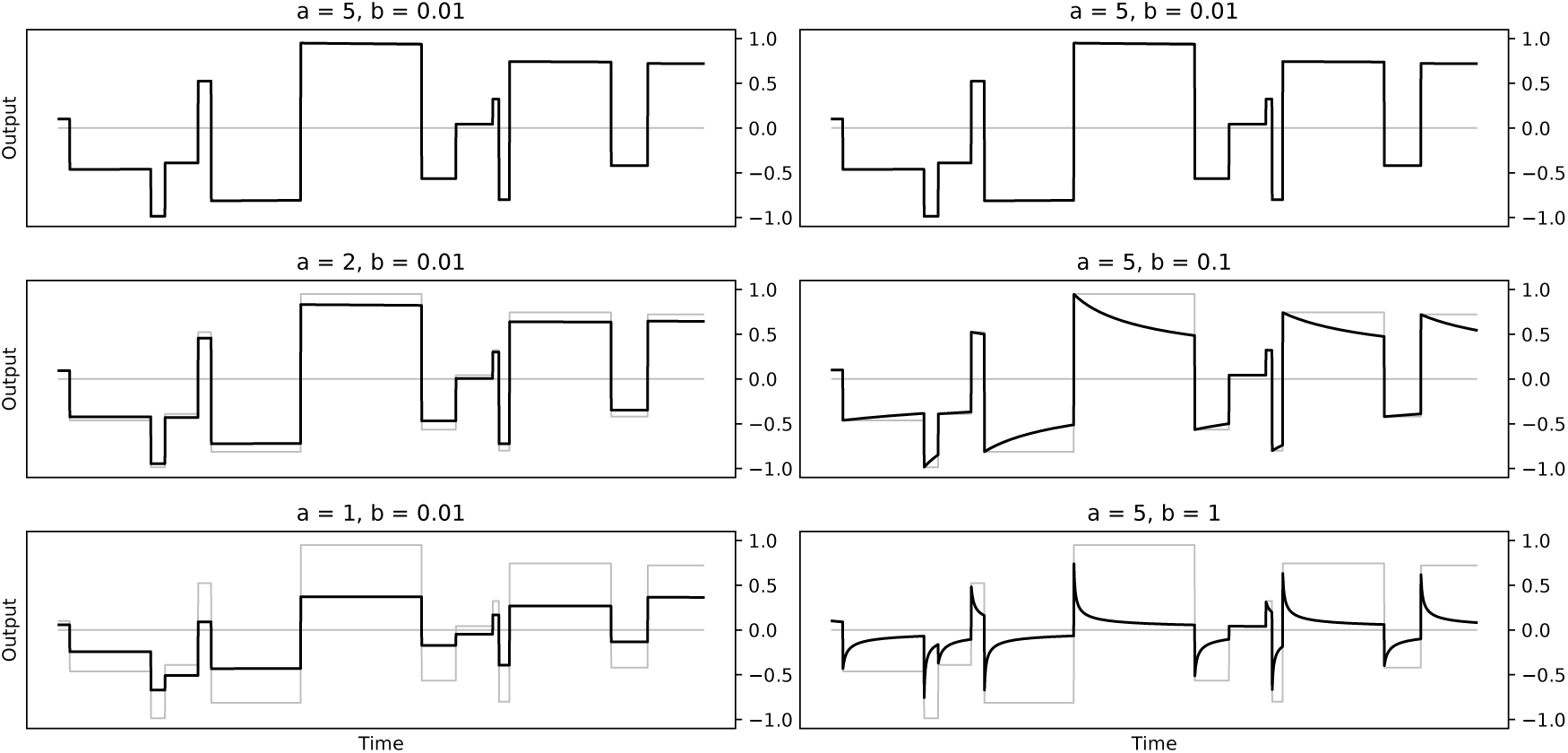
Influence of parameters *a* and *b* on the reduced model. The parameter *b* controls the amount of feedback fed to the model. If *b* is too high (*b* = 0.1, *b* = 1.0), the memory quickly fade to zero after a few timesteps. The parameter *a* controls the gate behavior and the ratio of the memorized and the new value that will constitute the new memorized value. If *a* is too low (*a* = 1, *a* = 2), there is a systematic undershoot.

### 5.2 Online learning

We tested online learning by using FORCE (Sussillo and Abbott, 2009) described by the following update equations:

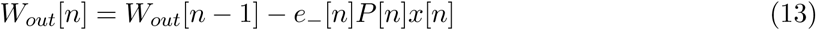

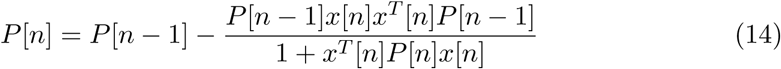

where *e*_*-*_[*n*] represents the error made at time *n* if there were no update of *W*_*out*_ and *P* [*n*] is a squared matrix computing an online inverse of *XX*^*T*^ *-αI, α* a regularization term and *P* initialized to 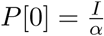.

In Figure 14 and 15, one can see that the behavior obtain with online learning is similar to offline learning.

**Figure 14:**
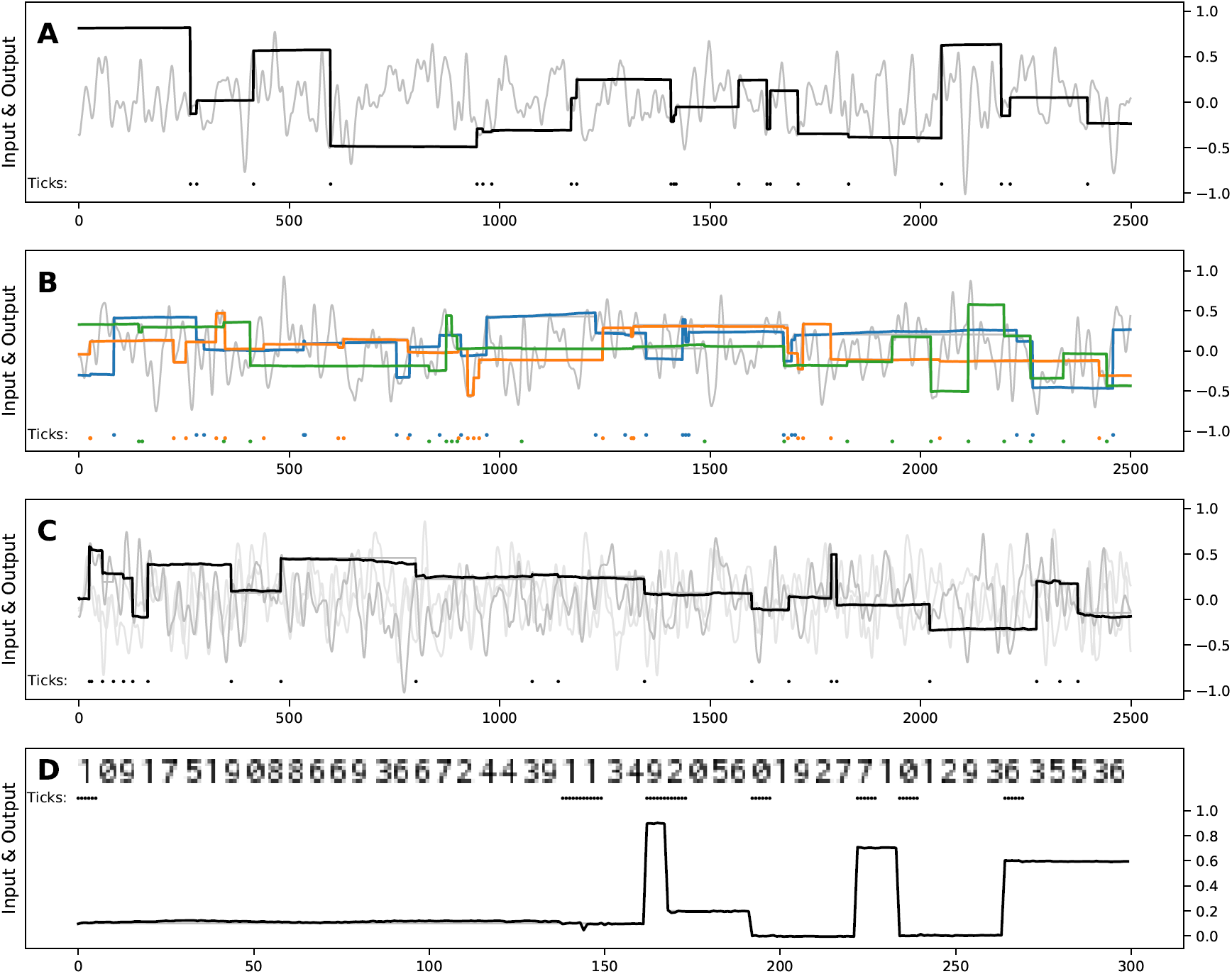
Performance of the reservoir model on working memory tasks with online FORCE learning. Same training/testing protocol than for Figure 9. The regularization parameter *α* has been fixed to 0.0001. The light gray line is the input signal and the thick black (or colored) one is the output of the model. Dots at the bottom represents the trigger signal (when to memorize a new value). For the digit task, the input containing the value to maintain is shown as a picture instead. **A** *1-value 1-gate scalar task* **B** *1-value 3-gate scalar task* **C** *3-value 1-gate scalar task* **D** *1-value 1-gate digit task*.

**Figure 15:**
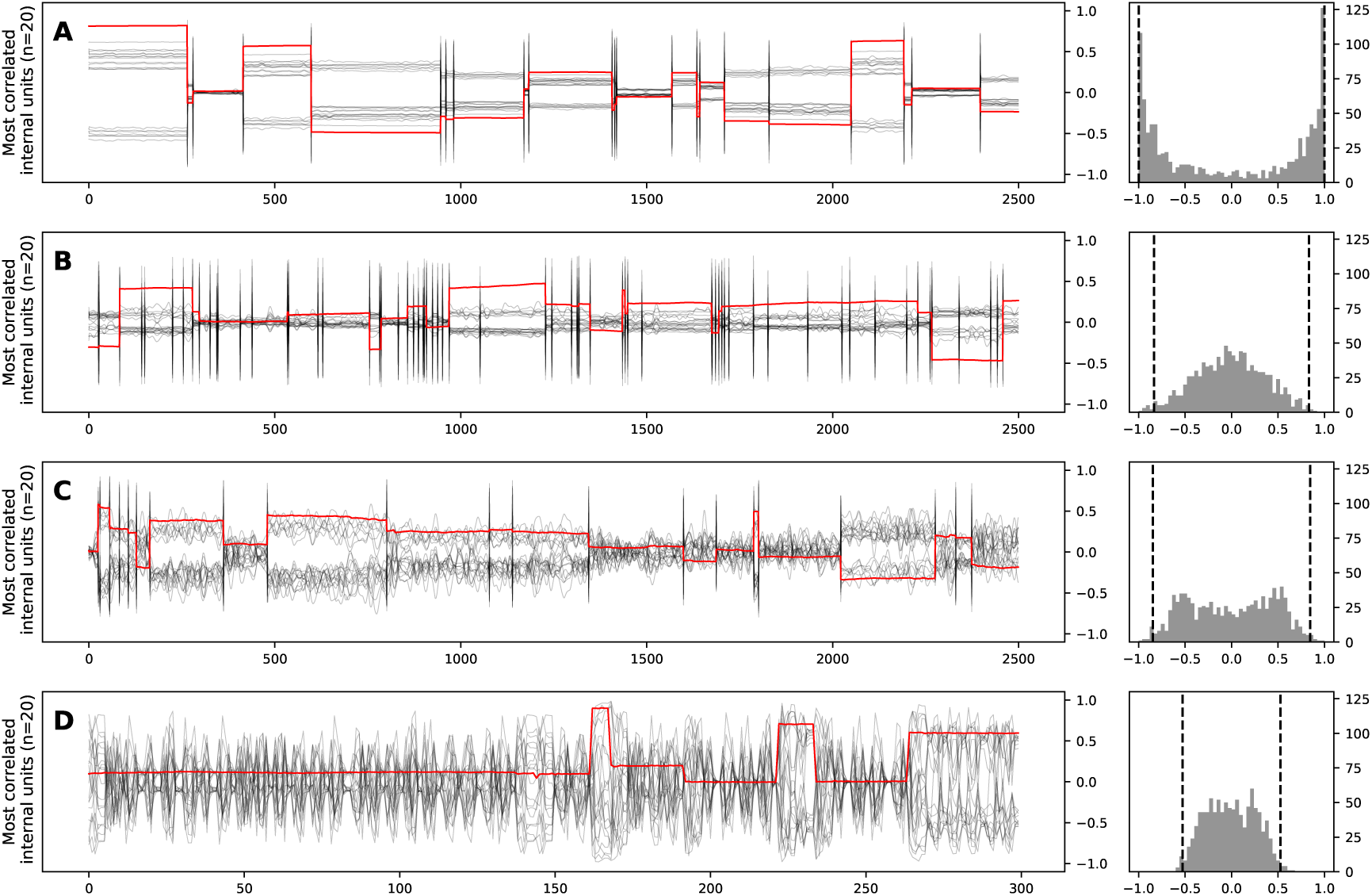
Most correlated reservoir units displaying various degrees of maintained activity with online FORCE learning. Same training/testing protocol than for Figure 9. The regularization parameter *α* has been fixed to 0.0001. (Left) in black the activities of the 20 neurons the most correlated with the output, in red the output of the model. (Right) histogram of the correlation between the neurons and the output; y-axis represents the number of neurons correlated with the output produced by the model within a correlation bin. Dashed lines indicate the 20th most correlated and most anti-correlatted neurons with the output. **A** *1-value 1-gate scalar task* **B** *1-value 3-gate scalar task* **C** *3-value 1-gate scalar task* **D** *1-value 1-gate digit task*. Most correlated reservoir units do not necessarily show clear sustained activity: we can see a degradation of sustained activities from **A** to **D** according to the different tasks. Thus, maintained activity is not compulsory to perform working memory tasks.

### 5.3 More on the segment attractor

In Figure 16 we performed a principal component analysis on the reservoir state obtained in Figure 8. We can note two interesting facts: (1) The activity evolves in a very low dimensional space, the first component explains more than 99% of the variance by itself, and (2) this component is linked to the memory maintained, linearly between −1 and 1 and non-linearly outside. With both combined together we can say that the segment attractor we built is actually a straight line.

**Figure 16:**
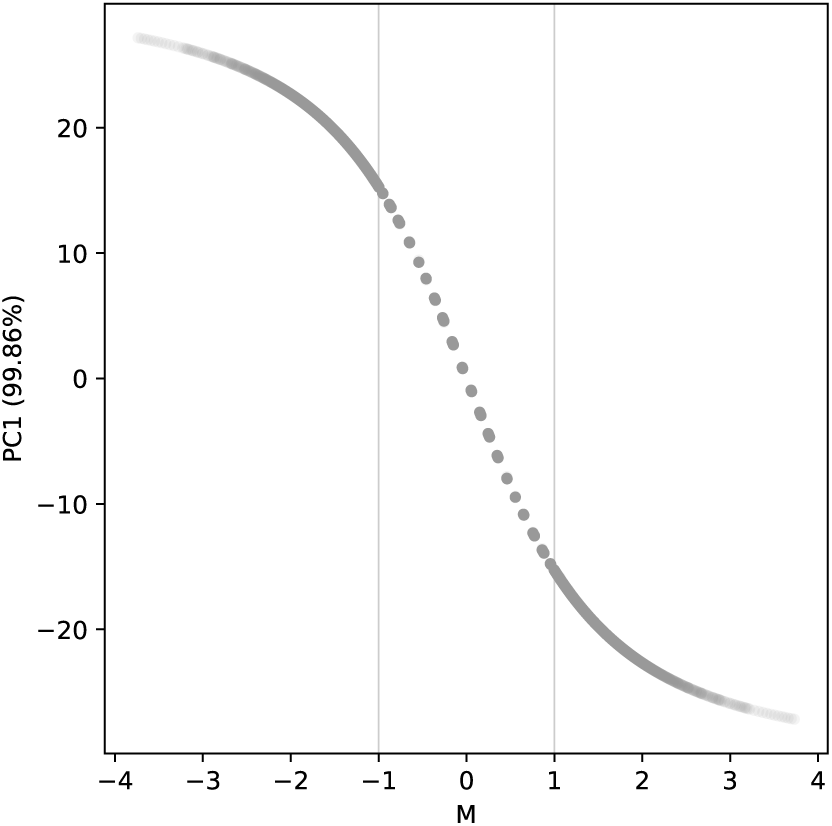
The line attractor. The reservoir state and output analyzed are the one of Figure 8. x-axis: output (memory). y-axis: first principal component of the reservoir state.

### 5.4 Influence of the spectral radius on the segment attractor

By watching how the segment attractor is evolving against the spectral radius, we can better explain its influence on the solving of the 1-value 1-gate task. The smaller the spectral radius the more precise is the approximation of the segment attractor. For a 0.1 spectral radius the approximate segment attractor seems continuous. The difference is quasi-null for starting values in [-1,+1] while for values outside this range, the difference corresponds to the difference with the nearest bound of the [-1,+1] segment. For a 0.5 spectral radius we clearly can see that the segment attractor is discretized. In the very first step the outputs concentrates around discretized value between −1 and 1 and is kept constant. In the extreme case for a 1.0 spectral radius it is discretized into two points corresponding to −1 and 1.

### 5.5 Killing neurons having a sustained activity

In Figure 18 we show how the neurons are correlated with the output (M), the value (V) and the trigger (T) for different feedback scaling. Because some irregularities come with triggers, we removed the trigger time steps while computing the correlation with the output and the value. We can note that the profile of neurons seems to change according to the feedback scaling. When it is high (1.0), there is mostly neurons following the output, when it is low (0.1) few neurons continue to follow the output while most of them vary according to the value they receive as input. To better identify the neurons which are following the output we looked at where were located the weights of these neurons in Figure 19. We can note that the most correlated neurons with the output are mainly the ones with relatively small input weights coming from the value. To go further with this idea we changed the sampling of the input weights to remove small values. In practice the sampling was done uniformly in [-1,-0.5]*∪*[0.5,1.0] instead. The behavior of this modified model is shown in Figure 20. We can note that the model is still able to perform the task relatively well, just a little worse than before, but there is no more neuron displaying a sustained activity. Interestingly, in Figure 19, the neurons the most correlated with the trigger displays an intringuing property, their output weights seems to be correlated to their feedback weights.

**Figure 17:**
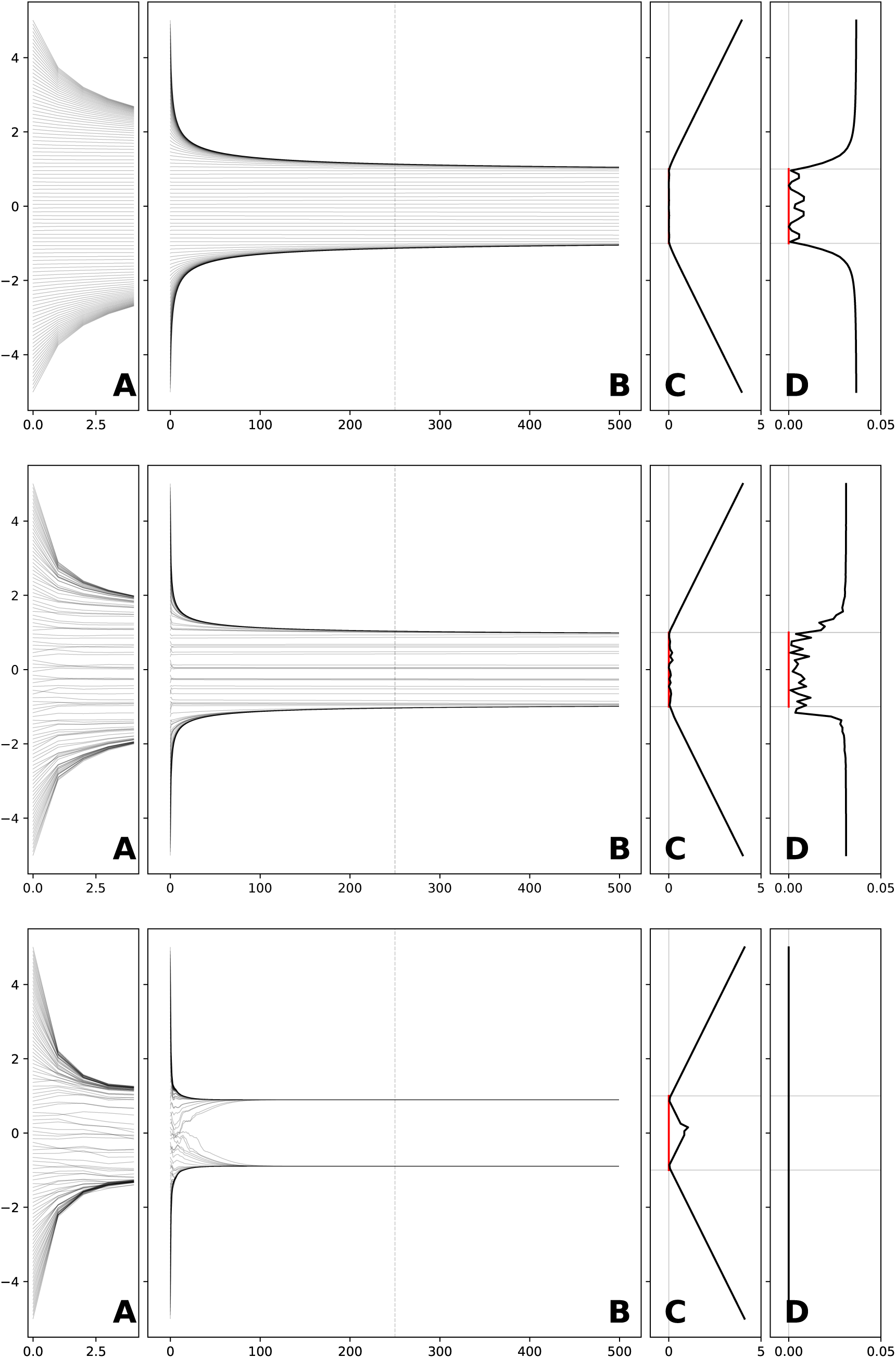
Influence of spectral radius on approximate line attractor. The same trained models were tested for 500 iterations without inputs, only starting with an initial trigger along with values linearly distributed between −5 and +5. First line: Spectral radius 0.1. Second line: Spectral radius 0.5. Third line: Spectral radius 1.0. **A** and **B** Output trajectories. Each line corresponds to one trajectory (output value) of the model. **A** is a zoom of **B** on the first few time steps. **C** Asolute difference in the output between initial time and and final time. **D** Maximal absolute difference in the neurons between intermediate (dashed line) and final time.

**Figure 18:**
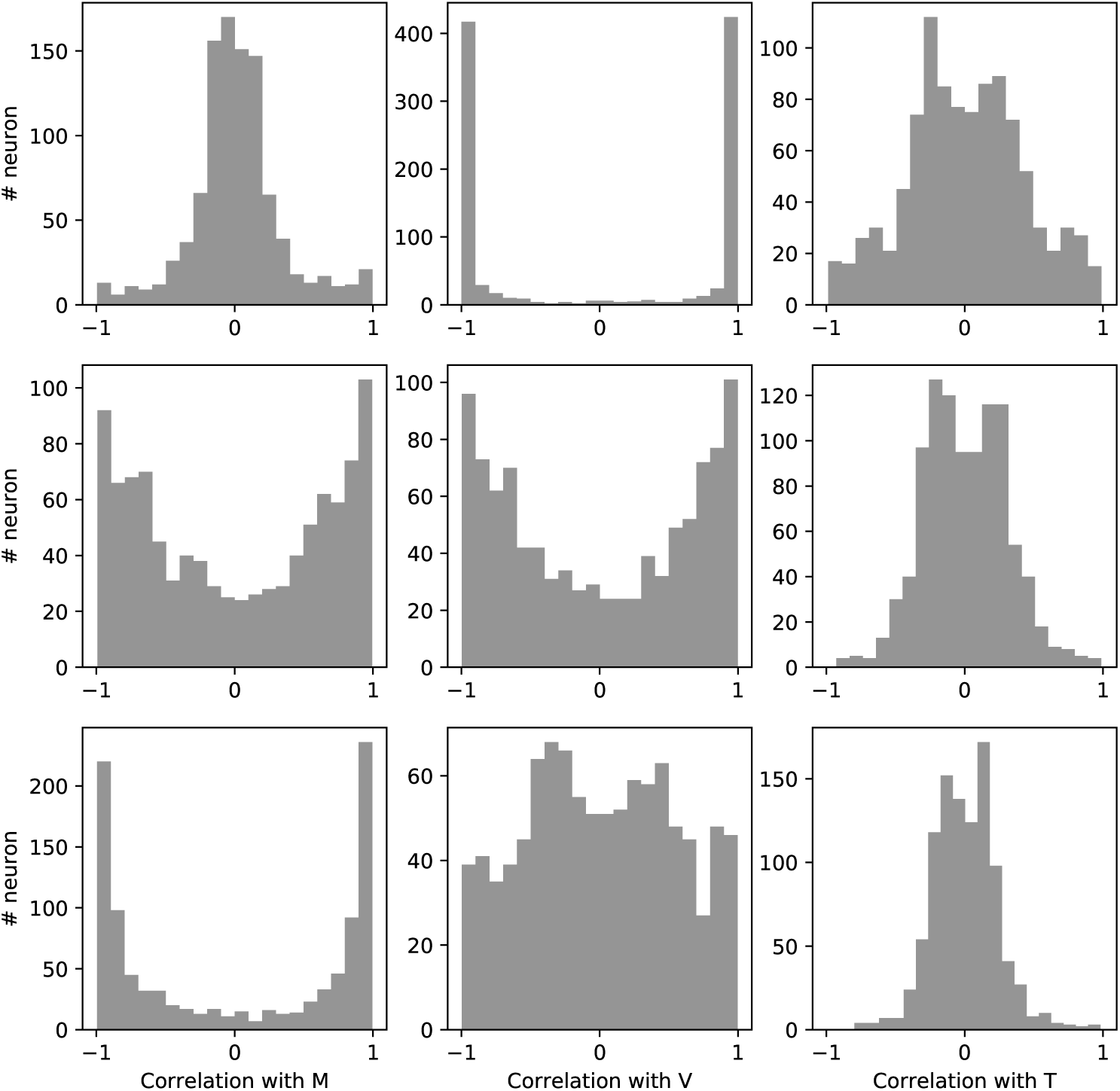
Influence of feedback scaling on maintenance of neurons. First line: Feedback scaling 0.1. Second line: Feedback scaling 0.5. Third line: Feedback scaling 1.0. First column: Correlation with M. Second column: Correlation with V. Third column: Correlation with T.

**Figure 19:**
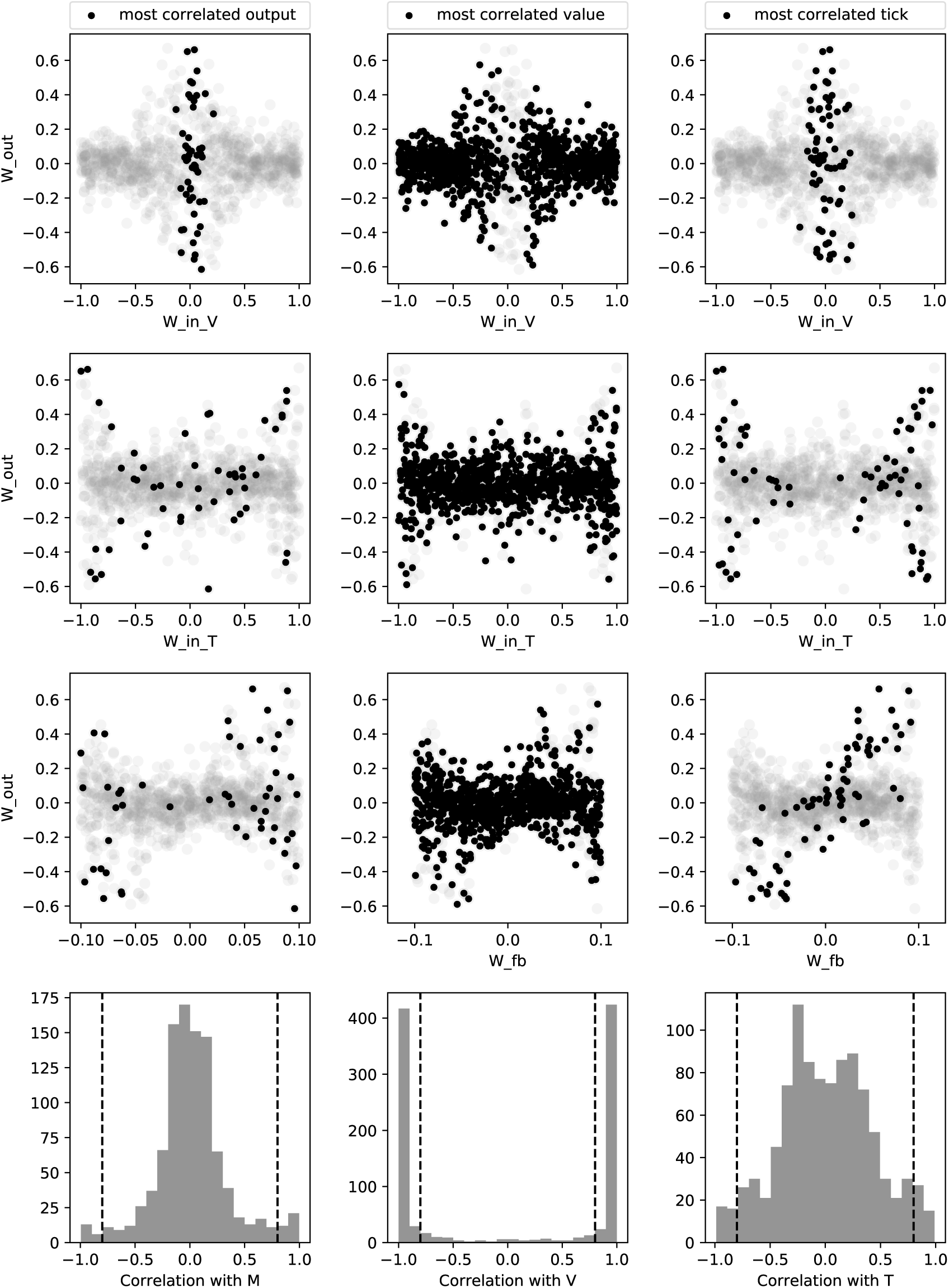
Link between weights and behavior of the neurons. The weights of the neurons the most correlated with either M, V or T are highlighted. First column: M. Second column: V. Third column: T. The three first lines show the link between the output weights and some other weights. First line: weights coming from V. Second line: weights coming from T. Third line: weights coming from M. The fourth line shows the correlation with the quantity used to highlight. The threshold used to decide which are the most correlated is shown in dotted black.

**Figure 20:**
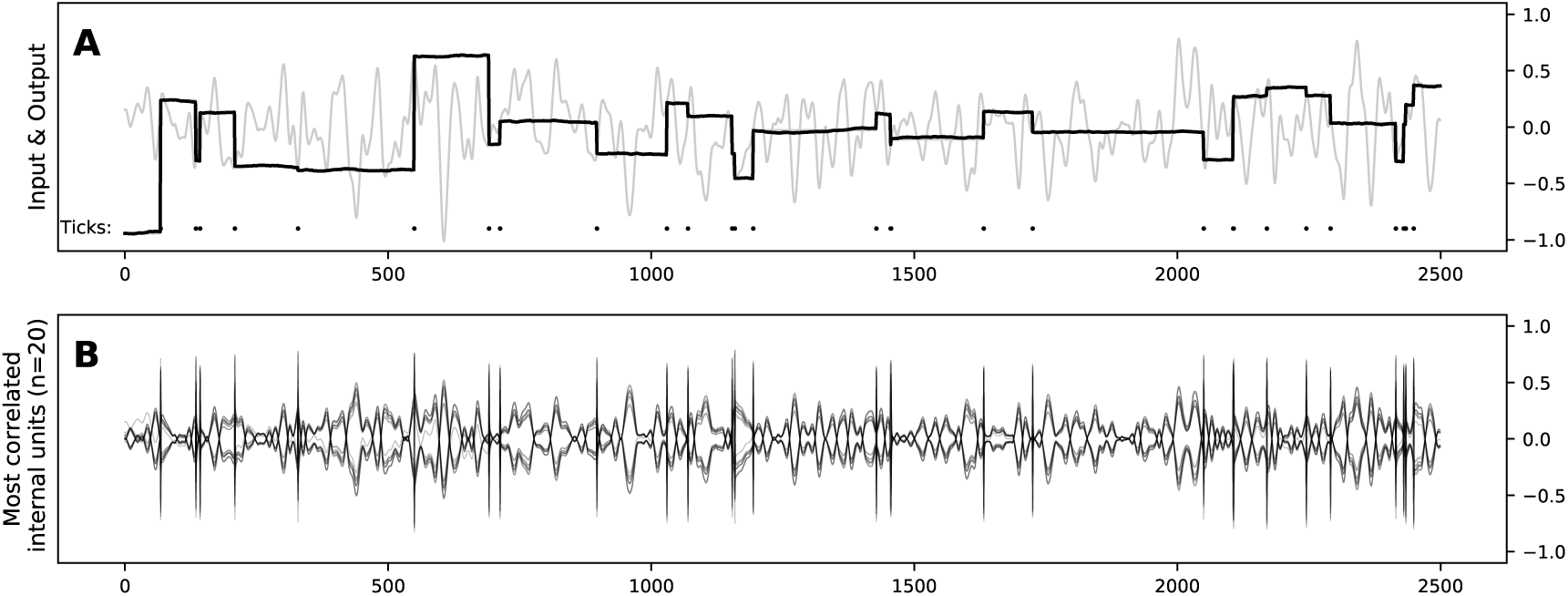
Performance of the model with high input value weights on the 1-gate task. **A** The light gray line is the input signal and the thick black one is the output of the model. Black dots at the bottom represents the trigger signal (when to memorize a new value). **B** Activity of 20 units that are the most correlated with the output.

### Footnotes

1 The Inconsolata font is available from https://www.levien.com/type/myfonts/inconsolata.html

2 We can actually find a similar LSTM variant in Greff et al., 2017 named *Coupled Input and Forget Gate*

3 Because a trigger input has a substantial influence on the reservoir states, we made the categorization by ignoring the time steps when there is a trigger.

4 Earlier formulations of very similar concepts can be found in (Jaeger, 2007).

5 For reservoirs this trade-off depends on the hyper-parameters (HP) chosen: some HP sets will give more memory, others more computational capacity (Legenstein and Maass, 2007).

